# A sensor of oxidative stress confers virulence via response memory in *Acinetobacter baumannii*

**DOI:** 10.64898/2026.04.08.717352

**Authors:** Hoan Van Ngo, Seung Hyeon Kim, Hongseok Ha, Sangwoo Kang, Donghyuk Shin, Matthias Gunzer, Kiwook Kim, Jinki Yeom

**Affiliations:** Department of Biomedical Science, College of Medicine, Seoul National University, Seoul, Korea; Department of Microbiology and Immunology, College of Medicine, Seoul National University, Seoul, Korea; Department of Pharmacology and Regenerative Medicine, University of Illinois College of Medicine, Chicago, IL, USA; Department of Systems Biology, College of Life Science and Biotechnology, Yonsei University, Seoul, Republic of Korea; Institute for Experimental Immunology and Imaging, University Hospital Essen, University of Duisburg-Essen, Essen, Germany; Leibniz-Institut für Analytische Wissenschaften - ISAS - e.V., Dortmund, Germany; University of Illinois Cancer Center, University of Illinois at Chicago, Chicago, USA; Cancer Research Institute, Seoul National University, Seoul, Republic of Korea

## Abstract

All living organisms must adapt to environmental stresses to survive. The two-component system (TCS) is a prevalent signal transduction mechanism to alter gene expression in response to stress in bacteria. We report that the PmrB sensor of the PmrA/PmrB TCS senses sublethal oxidative stress and encodes a response memory that promotes virulence in *Acinetobacter baumannii*. PmrB detects oxidative stress through histidine residues with a nickel (Ni^2+^) cofactor. Nickel oxidation (Ni^2+^) induces an allosteric effect by altering PmrB conformation, enabling PmrA activation. Activated PmrA induces antioxidant defense genes, including iron-sulfur cluster repair, ferritin, catalase, and peroxidase. Notably, PmrB remains activated after sublethal oxidative stress is removed, as pathogens encounter in the bloodstream and airways during infection. This response memory enables bacteria to respond strongly to subsequent lethal oxidative stress and antimicrobial peptides. In a murine model, disrupting oxidative sensing reduces virulence. In addition, PmrB-mediated response memory is essential for high virulence in clinically isolated multidrug-resistant *A. baumannii*. Our findings demonstrate how subinhibitory stress detection enables pathogens to achieve full virulence potential and resist stresses during infection.

## Introduction

All living organisms must adapt to environmental stresses to survive and proliferate. These stresses often serve as signals that stimulate the organism to adjust its physiology (Storz *et al*, 2011). In bacteria, the most common mechanism for stress responses is the two-component signal transduction regulatory system (TCS) (Groisman & Mouslim, 2006). A TCS generally comprises a sensor kinase that alters the phosphorylation state of a transcriptional regulator in response to a particular signal (Lazar & Tabor, 2021; Stock *et al*, 2000; Zschiedrich *et al*, 2016). In pathogenic bacteria, TCS governs global transcriptional regulation in response to signals during infections (Lazar & Tabor, 2021; Stock *et al*., 2000; Zschiedrich *et al*., 2016). Notably, pathogenetic bacteria encounter sublethal oxidative stress in the host’s bloodstream and airways during infection (Forman *et al*, 2016; Padron *et al*, 2023). It is poorly known how sublethal stress regulates TCS response in the host. In this study, we investigate that sublethal stress establishes response memory in TCS that significantly increases the virulence of bacterial pathogens during infections.

*Acinetobacter baumannii* is a Gram-negative pathogenic bacterium that predominantly causes nosocomial infections (Dijkshoorn *et al*, 2007). It can infect various parts of the human body, including the blood, lungs, urinary tract, and wounds (Ayoub Moubareck & Hammoudi Halat, 2020; Weiner-Lastinger *et al*, 2020). For successful infection, *A. baumannii* must defend against environmental stresses produced by the host immune system, including oxidative stress and antimicrobial peptides (Jones *et al*, 2017; Juttukonda *et al*, 2019; Qiu *et al*, 2009). Importantly, *A. baumannii* poses a global threat due to the increasing multidrug-resistant isolates from clinical settings (Levy-Blitchtein *et al*, 2018; Nasr, 2020; Talbot *et al*, 2006). Antimicrobial peptides such as colistin are a last-resort drug for treating multidrug-resistant *A. baumannii* infections (Gurjar, 2015; Novovic & Jovcic, 2023). However, it is presently unclear how *A. baumannii* resists antimicrobial peptides in the host during infections. This study demonstrates that *A. baumannii* resists high concentrations of antimicrobial peptides through cross-protection, mediated by the PmrA/PmrB system’s response memory, primed by sublethal oxidative stress.

Cells use respiration to obtain energy, but aerobic respiration can produce reactive oxygen species (ROS), such as superoxide (O₂·) and hydrogen peroxide (H₂O₂), which are toxic byproducts that can damage DNA, proteins, and lipids (Bonora *et al*, 2012; Imlay, 2008). Subsequently, H₂O₂ reacts rapidly with free ferrous iron (Fe²⁺), generating highly toxic hydroxyl radicals (HO·) through the Fenton reaction (Imlay *et al*, 1988). To survive under oxidative stress, living cells must protect their macromolecules from ROS-induced damage. For instance, in many bacteria, the transcriptional regulator OxyR and the sigma factor RpoS activate genes encoding enzymes that degrade H₂O₂ (Chiang & Schellhorn, 2012). Additionally, the assembly systems of iron-sulfur (Fe-S) clusters, including Nif, Isc, and Suf, can help to prevent and repair Fe-S cluster from ROS damage (Angelini *et al*, 2008; Jang & Imlay, 2010; Lee *et al*, 2004; Lill & Muhlenhoff, 2005; Nachin *et al*, 2001; Nachin *et al*, 2003; Yeo *et al*, 2006). Intriguingly, *A. baumannii* does not encode homologs of RpoS, Nif, or Suf proteins (Appendix dataset S1), suggesting that other regulatory systems may be involved in responses to and defense against oxidative stress. Here, it is revealed that a PmrA TCS plays a role in oxidative stress defense, which fully activates virulence in *A. baumannii*.

In this study, we demonstrate that the *A. baumannii* sensor PmrB from the PmrA/PmrB TCS system directly detects peroxides through histidine residues with a nickel cofactor. Oxidation of the nickel cofactor by oxidative stress in PmrB activates the transcriptional regulator PmrA, which then regulates the expression of genes involved in oxidative stress defense. Notably, we discovered a new signal-priming mechanism in which exposure to sublethal concentrations of oxidative stress forms response memory of the PmrA/PmrB system, conferring robust and strong cross-protection against lethal concentrations of both oxidative stress and antimicrobial peptides. This mechanism is required for full virulence activation of *A. baumannii* in the host. Moreover, the histidine box, which binds the nickel cofactor, is a determinant of hypervirulence in clinically isolated multidrug-resistant *A. baumannii*. This represents a previously unrecognized strategy by which bacterial pathogens prepare for hostile host environments during infections by sensing subinhibitory stress.

## Results

### The PmrA/PmrB is required for defense against oxidative stress in *A. baumannii*

Eukaryotic hosts use oxidative stress to defend against pathogens within innate immune cells (Herb & Schramm, 2021; Nguyen *et al*, 2017; Segal, 2005). Pathogens also encounter oxidative stress from fluid flow in the host’s bloodstream and airways during infection (Forman *et al*., 2016; Padron *et al*., 2023). As a result, it is essential that pathogenic bacteria have evolved mechanisms to withstand oxidative stress and survive and proliferate in the host. Interestingly, several bacterial species, including *A. baumannii*, lack homologs of *rpoS*, *nif*, and *suf,* which are required for defense against oxidative stress (Appendix dataset S1). These pathogenic bacteria must still prepare efficient oxidative stress defense to survive during infections. Thus, we reasoned that other regulatory systems, such as the two-component signal transduction regulatory systems (TCSs), may help defend against oxidative stress.

While the PmrA/PmrB TCS confers antimicrobial peptide resistance in *Salmonella*, *A. baumannii* PmrB has a distinct structure (Fig. 1A) and only 28.57% sequence similarity with *Salmonella* PmrB (Appendix Fig. S1). Also, *A. baumannii* PmrB forms a unique clade in the phylogenetic tree (Appendix Fig. S2), and its periplasmic domain, which is typical signal sensing domain, has a barrel structure, while *Salmonella*’s has a coil structure (Fig. 1A). Barrel structures be often involved in oxidative stress sensing in other enzymes (Perez-Dominguez *et al*, 2021; Tala *et al*, 2018). Given these differences and the lack of conventional oxidative stress defenses in *A. baumannii*, we hypothesized that the *A. baumannii* PmrA/PmrB system may be involved in defense against oxidative stress.

**Figure 1.**
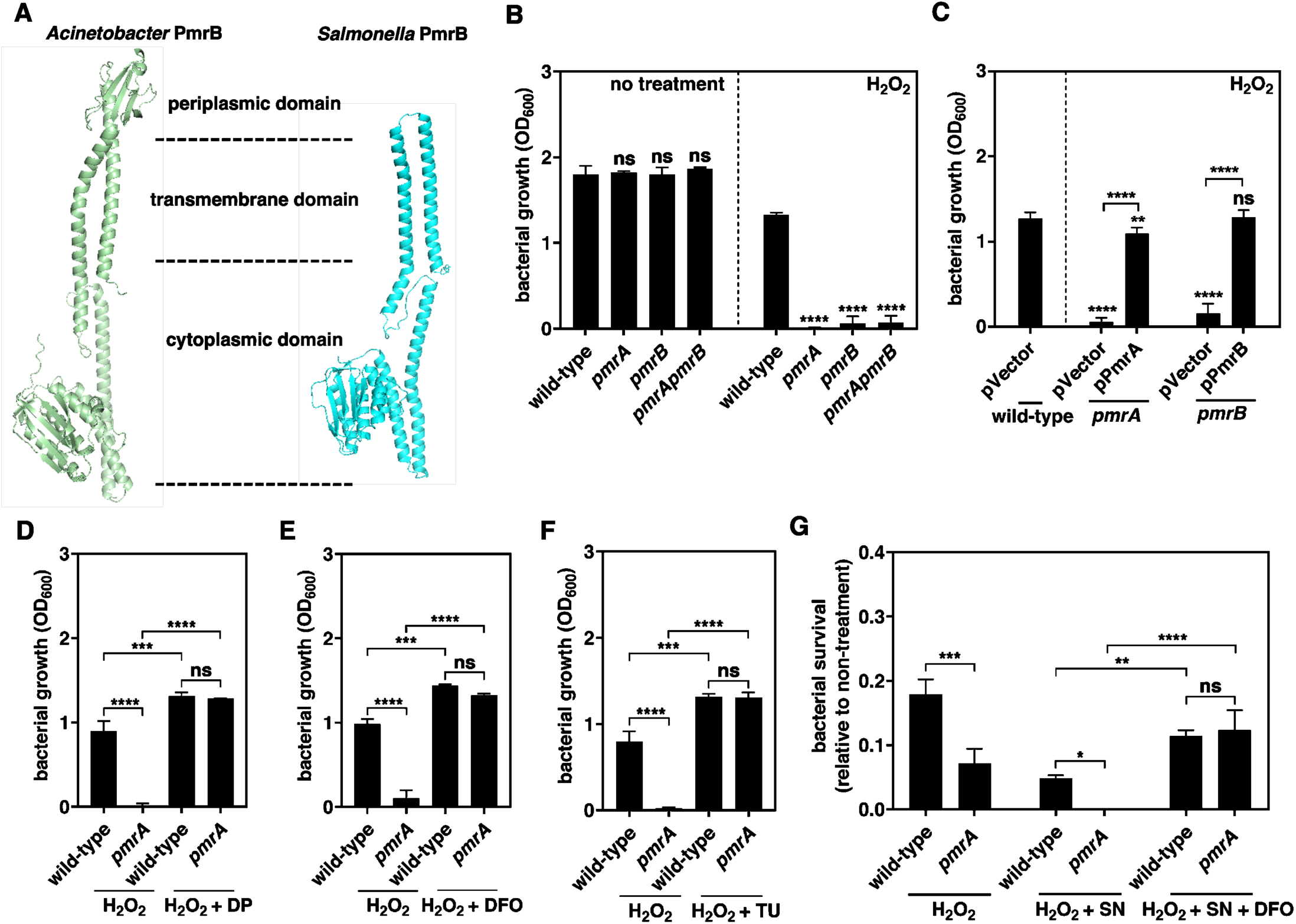
The PmrA/PmrB system is important to defense against oxidative stress in *A. baumannii.* (A) Predicted structures of PmrB in *A. baumannii* and *Salmonella* by Alphafold 2 program. (B) Growth of *A. baumannii* 17978 wild-type, *pmrA*, *pmrB*, and *pmrA pmrB* in LB media alone (no treatment) or LB media containing 2 mM H_2_O_2_ at 37°C after 20 h. (C) Growth of *A. baumannii* 17978 wild-type harboring the plasmid vector (pWH1266), *pmrA* harboring the plasmid vector or the *pmrA*-expressing plasmid (pPmrA), and *pmrB* harboring the plasmid vector or the *pmrB*-expressing plasmid (pPmrB) in LB media containing 2 mM H_2_O_2_ at 37°C after 20 h. (D-F) Growth of *A. baumannii* 17978 wild-type and *pmrA* in LB media supplemented with 2 mM H_2_O_2_ alone or in combination with 100 uM 2,2’-dipyridyl (DP) (D), 100 uM deferoxamine (DFO) (E), or 10 mM thiourea (TU) (F) at 37°C after 20 h. (G) Relative survival of *A. baumannii* 17978 wild-type and *pmrA* after treatment with 10 mM H_2_O_2_ or in combination with 40 µg/mL streptonigrin (SN) and 100 µM deferoxamine (DFO). Data are presented as means ± SD from n = 3. *P* values were determined using one-way ANOVA with Dunnett’s multiple comparision test (B and C) or two-way ANOVA with Tukey’s multiple comparision test (D, E, F and G). *P < 0.05, **P < 0.01, ***P < 0.001 and ****P < 0.0001, ns; not significant. Exact P values in Appendix Table S1.

Multiple lines of evidence support that the PmrA/PmrB system is required for oxidative stress defense in *A. baumannii*. First, exposure to hydrogen peroxide (H_2_O_2_), an oxidative stressor, led to complete growth defects in mutant strains lacking *pmrA*, *pmrB*, or both. In contrast, the wild-type strain can grow under oxidative stress, and mutant strains of *pmrA* and/or *pmrB* genes did not have any growth defect without oxidative stress (Fig. 1B). Second, plasmids harboring either *pmrA* or *pmrB* corrected the growth defect in the *pmrA* or *pmrB* mutants under H_2_O_2_ treatment, whereas the empty vector did not (Fig. 1C). Third, inactivation of *pmrA* or *pmrB* impaired bacterial survival under treatment of spermine NONOate (SPE), a nitric oxide generator (Appendix Fig. S3A). Fourth, plasmids harboring either *pmrA* or *pmrB* corrected survival defects in the mutants under SPE treatment, whereas the empty vector did not (Appendix Fig. S3B). Lastly, although OxyR is well known as a transcriptional regulator of the oxidative stress response in many bacteria (Zheng *et al*, 1998), *oxyR* inactivation did not affect bacterial survival upon H_2_O_2_ exposure in *A. baumannii,* unlike *pmrA* inactivation (Appendix Fig. S3C), consistent with previous findings (Juttukonda *et al*., 2019). Therefore, the two-component regulatory system PmrA/PmrB is required for oxidative stress defense in *A. baumannii*.

### The PmrA/PmrB defends against the Fenton reaction-mediated oxidative stress

In living cells, the Fenton reaction causes strong oxidative stress by generating highly toxic hydroxyl radicals (HO·) in the cytoplasm, driven by peroxide and free ferrous iron (Imlay *et al*., 1988). To determine whether the detrimental effects of H_2_O_2_ originated from the Fenton reaction, we treated bacteria with iron chelators or hydroxyl radical scavengers to inhibit the Fenton reaction-mediated oxidative stress.

The ferrous iron chelator 2,2’-dipyridyl (DP) restored bacterial growth defect of the *pmrA* mutant under H_2_O_2_ treatment compared to wild-type levels (Fig. 1D). Similarly, deferoxamine (DFO), which chelates both ferric and ferrous iron, enhanced bacterial growth of both wild-type and *pmrA* mutant strains under H_2_O_2_ treatment (Fig. 1E). Additionally, thiourea (TU) treatment, a hydroxyl radical scavenger, recovered growth defect under H_2_O_2_ treatment (Fig. 1F). Additionally, streptonigrin (SN), which kills cells dependent on cytoplasmic free ferrous iron levels (Gupta & Imlay, 2023), combined with H_2_O_2_ promotes more bacteria killing than only H_2_O_2_ treatment (Fig. 1G). Also, *pmrA* inactivated *A. baumannii* could not survival at all under SN and H_2_O_2_ treatment (Fig. 1G). By contrast, the iron chelator DFO improved survival and removed the survival difference in both wild-type and *pmrA* mutant strains (Fig. 1G). In addition, TU treatment restored the survival defect induced by SN and H_2_O_2_ in wild-type and *pmrA* mutant strains (Appendix Fig. S3D). Collectively, these findings show that the PmrA/PmrB system is essential for oxidative stress defense in *A. baumannii,* mainly by reducing the Fenton reaction.

### The PmrA/PmrB promotes gene expression involved in oxidative stress defense

Next, we investigated how the PmrA/PmrB system helps bacteria cope with oxidative stress. As the Isc system is the sole Fe-S cluster assembly machinery in *A. baumannii* (Appendix dataset S1), and free cytoplasmic iron promotes oxidative stress (Imlay *et al*., 1988) (Fig. 1D and E). In addition, FtnA protein is the main scavenger of free ferrous iron (Fe²⁺) in bacteria, preventing Fenton reactions by reducing free cytoplasmic iron-induced oxidative stress (Abdul-Tehrani *et al*, 1999; Faulkner & Helmann, 2011; Matveeva *et al*, 2025). Given this, we hypothesized that PmrA may promote the expression of the Isc system to repair Fe-S cluster damage caused by ROS and the free-iron scavenger FtnA, thereby reducing Fenton reactions.

Multiple lines of evidence support the idea that PmrA directly activates the expression of genes involved in Fe-S cluster assembly and a free-iron scavenger under oxidative stress conditions. First, a putative PmrA binding site (5’-TTTAAT N3 TTTAAT-3’) is located upstream of the *hscB* (Wosten & Groisman, 1999). This is the first gene of the *hscB-hscA-fdx* operon (Fig. 2A), which belongs to the Isc system in *A. baumannii*. This operon might be involved in the repair of the iron-sulfur cluster in *A. baumannii*. Second, H_2_O_2_ induced a ∼6-fold increase in *lux* expression in bacteria harboring the *hscB-hscA-fdx* promoter region cloned into a luciferase reporter system. Inactivation of *pmrA* significantly reduced this activity under H_2_O_2_. By contrast, colistin, which is a known stress molecule and can activate the PmrA/PmrB system, did not induce the *hscB*-*hscA*-*fdx* expression (Fig. 2B). Third, purified PmrA protein shifted a DNA fragment harboring the *hscB-hscA-fdx* promoter region in electrophoretic mobility shift assays (EMSA) (Fig. 2C). By contrast, excess unlabeled promoter DNA competes out binding between PmrA and labeled *hscB-hscA-fdx* promoter DNA (Fig. 2C). Fourth, a putative PmrA binding site is also located upstream of *ftnA* (Appendix Fig. S4A). Fifth, *ftnA* gene expression was strongly upregulated when bacteria were exposed to H_2_O_2_, but not colistin (Appendix Fig. S4B). Finally, purified PmrA shifted a DNA fragment containing the *ftnA* promoter in protein-DNA binding assays (Appendix Fig. S4C). Excess unlabeled *ftnA* promoter DNA competed out labeled *ftnA* promoter to interact with PmrA (Appendix Fig. S4C). Together, these data indicate that PmrA directly activates the iron-sulfur cluster repair systems and the free-iron scavenger FtnA in response to oxidative stress, which enables bacteria to reduce cytoplasmic free ferrous iron.

**Figure 2.**
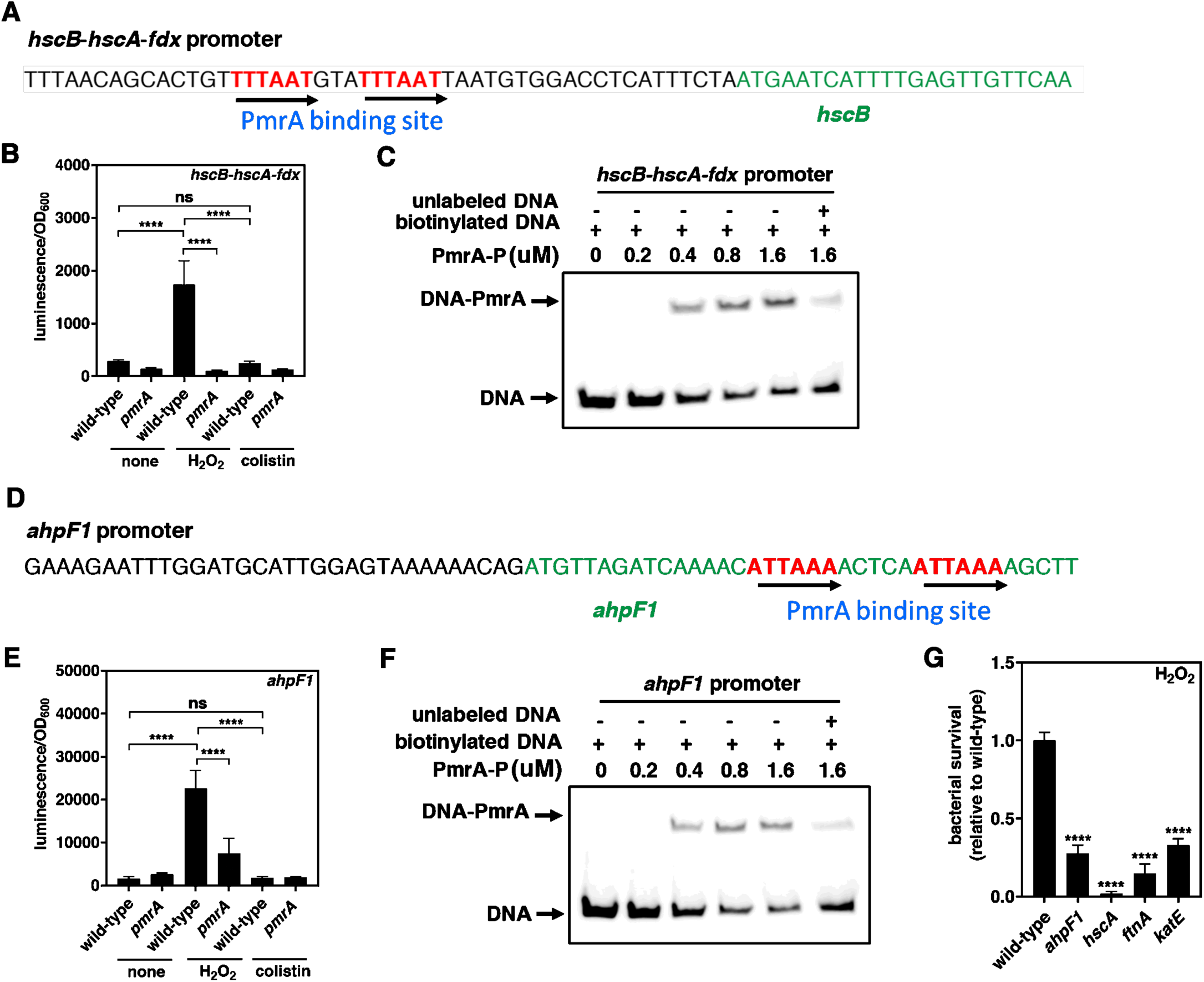
The response regulator directly binds to *hscB*-*hscA*-*fdx* and *ahpF1* promoters and regulate its expression in response to oxidative stress. (A) The schematic localization of the putative PmrA binding sites (red color) in the *hscB-hscA-fdx* promoter and open reading frame for *hscB* gene (green color). The binding sites are TTAAT N3 TTTAAT for *hscB-hscA-fdx* promoter. (B) Luminescence emission of *A. baumannii* 17978 carrying the luciferase reporter plasmid pLPV1Z fused with *hscB-hscA-fdx* promoter at 2 h in the presence of 10 mM H_2_O_2_ or 1 µg/mL colistin. (C) Electrophoretic mobility shift assay (EMSA) validates that the PmrA interaction with the *hscB-hscA-fdx* promoter. (D) The schematic localization of the putative PmrA binding sites (red color) in the *ahpF1* promoter and open reading frame for *ahpF1* gene (green color). The binding sites are ATTAAA N5 ATTAAA for *ahpF1* promoter. (E) Luminescence emission of *A. baumannii* 17978 carrying the luciferase reporter plasmid pLPV1Z fused with *ahpF1* promoter at 2 h in the presence of 10 mM H_2_O_2_ or 1 µg/mL colistin. (*F*) EMSA validates that the PmrA interaction with the *ahpF1* promoter. (*G*) Relative survival of *A. baumannii* 17978 wild-type and *ahpF1*, *hscA*, *ftnA*, and *katE* mutants after treatment with 10 mM H_2_O_2_. Data are presented as means ± SD from n = 3. *P* values were determined using two-way ANOVA with Tukey’s multiple comparision test (B and E) or one-way ANOVA with Dunnett’s multiple comparision test (G). ****P < 0.0001, ns; not significant. Exact P values in Appendix Table S1.

Furthermore, we tested whether PmrA directly activates the *ahpF1* and *katE* genes, which encode the major peroxidase and catalase enzymes to degrade H_2_O_2_ in *A. baumannii* (Juttukonda *et al*., 2019). A putative PmrA box was identified near the translation start site of *ahpF1* and *katE* (Fig. 2D and Appendix Fig. S4D). Luciferase reporter assays revealed strong upregulation of *ahpF1* and *katE* expressions in response to H_2_O_2_, which was significantly lower in the *pmrA* mutant (Fig. 2E and Appendix Fig. S4E). By contrast, colistin did not promote *ahpF1* and *katE* expression (Fig. 2E and Appendix Fig. S4E). Also, PmrA directly binds to the *ahpF1* and *katE* promoters (Fig. 2F and Appendix Fig. S4F). Excess unlabeled *ahpF1* and *katE* promoter DNAs competed out labeled promoter DNAs to bind to purified PmrA (Fig. 2F and Appendix Fig. S4F). By contrast, gene expression of *katG*, which is another catalase in *A. baumannii*, was not activated by PmrA under oxidative stress (Appendix Fig. S4G), and PmrA did not bind to the *katG* promoter region (Appendix Fig. S4H). Importantly, inactivation of *ahpF1, hscA, ftnA*, or *katE* caused a significant reduction in the survival of bacteria exposed to H_2_O_2_ (Fig. 2G).

Moreover, OxyR is a transcriptional activator of oxidative stress responses in many bacteria and functions as a cytoplasmic H_2_O_2_ sensor that activates gene expression by forming an intermolecular disulfide bond (Zheng *et al*., 1998). To test whether the PmrA/PmrB system is related to the OxyR regulator, we measured *ahpF1*expression in *oxyR* or *pmrA* mutant strains. As expected, OxyR and PmrA can activate gene expression of ahpF1 under oxidative stress, inactivation of *oxyR* or *pmrA* significantly reduced expression of the *ahpF1* gene (Appendix Fig. S4J). Interestingly, *A. baumannii* OxyR repressed *ahpF1* expression under normal conditions (Appendix Fig. S4J), consistent with a previous finding (Juttukonda *et al*., 2019). By contrast, PmrA did not regulate *ahpF1* expression in the absence of oxidative stress (Appendix Fig. S4J). Thus, OxyR and PmrA are independent pathways to regulate oxidative stress defense system in *A. baumannii*. Taken together, PmrA directly activates the expression of the iron-sulfur repair system, the free iron scavenger, peroxidase, and catalase, which are key components of the oxidative stress defense in *A. baumannii*.

### PmrB senses oxidative stress via histidine residues in the periplasmic domain

To defend against oxidative stress, the PmrB sensor of the PmrA/PmrB system must sense oxidative stress molecules. To investigate how the PmrB recognizes oxidative stress molecules in *A. baumannii,* we identified a critical domain for oxidative stress sensing in the PmrB sensor. PmrB is a multidomain transmembrane protein with an N-terminal cytoplasmic domain, two transmembrane domains, a periplasmic domain, a histidine kinase domain, and an HATPase-c domain (Fig. 3A). Notably, the periplasmic domain of *A. baumannii* PmrB has significantly different structure from that of *Salmonella* PmrB (Fig. 1A). In addition, the *Salmonella* PmrA/PmrB TCS did not sense and respond to oxidative stress (Wosten *et al*, 2000). Based on these findings, we hypothesized that the periplasmic domain of PmrB may be necessary for sensing oxidative stress in *A. baumannii*.

**Figure 3.**
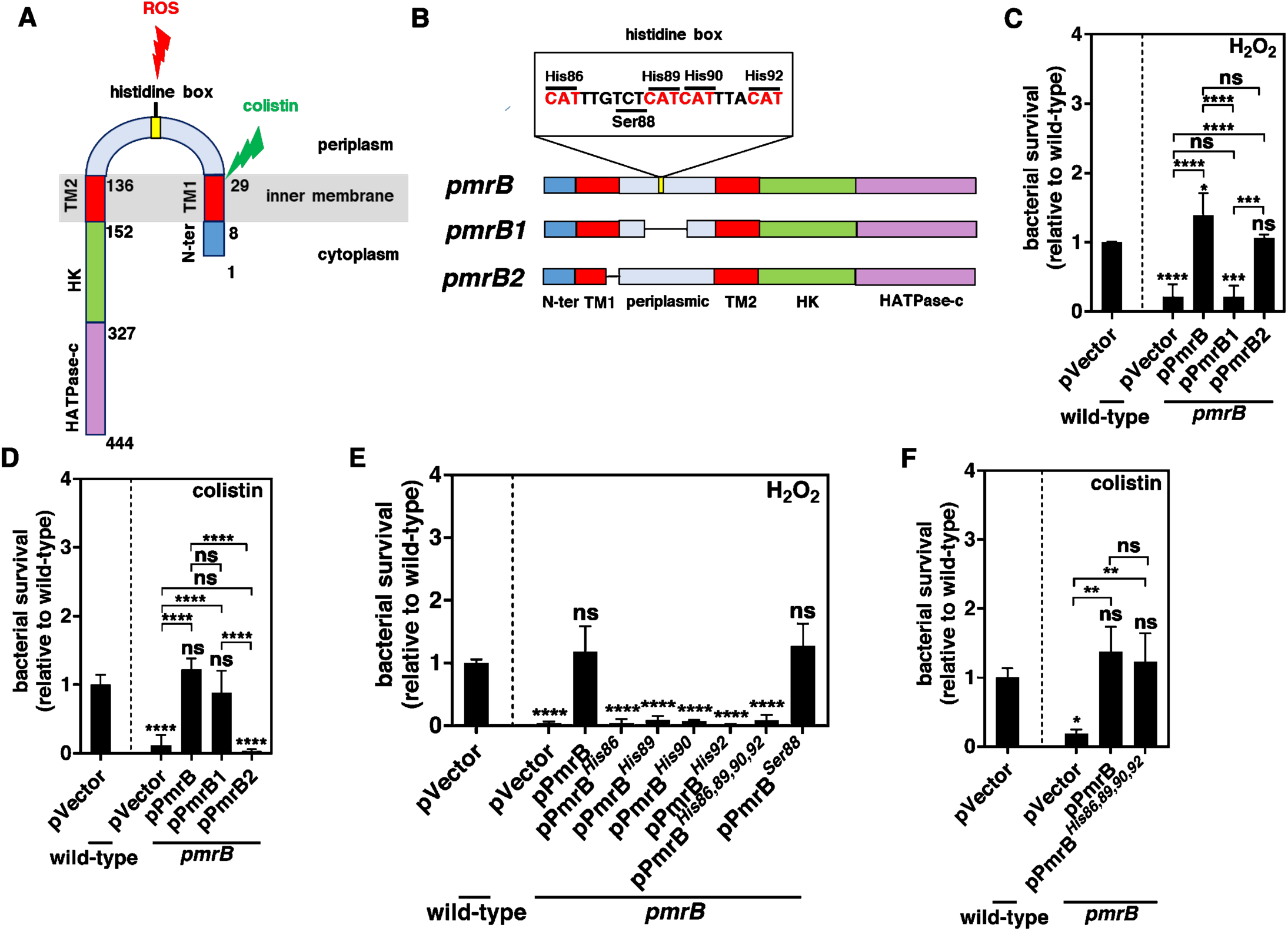
Histidine residues on the periplasmic domain of PmrB is critical for the ability of PmrB to sense oxidative stress. (A) Schematic representation of histidine kinase PmrB. PmrB domains: N-ter, N-terminal cytoplasmic domain (aa 1-8); TM1, first transmembrane domain (aa 9-29); periplasmic domain that contains histidine box (aa 30-136); TM2, second transmembrane domain (aa 137-152); HK, histidine kinase (aa 153-327), and HATPase-c, histidine kinase-like ATPases (aa 328-444). Red thunder indicates site for ROS sensing while green thunder indicate site for colistin sensing. (B) Schematic representation of full-length PmrB and its variants (PmrB1 lacking aa 70-130 and PmrB2 lacking aa 23-29). Full-length PmrB consists of a histidine box of 4 histidine residues (His86, His89, His90, and His92) on the periplasmic domain, whereas PmrB1 lacks aa 70-130 (no histidine box). (C, D) Relative survival of *A. baumannii* 17978 wild-type harboring the plasmid vector (pWH1266) and *pmrB* harboring the plasmid vector, the *pmrB*-expressing plasmid (pPmrB), the *pmrB1*-expressing plasmid (pPmrB1), or the *pmrB2*-expressing plasmid (pPmrB2) after treatment with 10 mM H_2_O_2_ (C) or 1 µg/mL colistin (D). (E) Relative survival of *A. baumannii* 17978 wild-type harboring the plasmid vector (pWH1266) and *pmrB* harboring the plasmid vector, the *pmrB*-expressing plasmid (pPmrB), or the plasmid expressing *pmrB* mutated in histidine residues (pPmrB*^His86^*, pPmrB*^His89^*, pPmrB*^His90^*; pPmrB*^His92^* or pPmrB*^His86,89,90,92^*) or serine residue (pPmrB*^Ser88^*) after treatment with 10 mM H_2_O_2_. (F) Relative survival of *A. baumannii* 17978 wild-type harboring the plasmid vector (pWH1266) and *pmrB* harboring the plasmid vector, the *pmrB*-expressing plasmid (pPmrB), or the plasmid expressing *pmrB* mutated in histidine residues (pPmrB*^His86,89,90,92^*) after treatment with 1 µg/mL colistin. Data are presented as means ± SD from n = 3. *P* values were determined using one-way ANOVA with Tukey’s multiple comparision test (C, D, and F) or Dunnett’s multiple comparitions test ©. *P < 0.05, **P < 0.01, ***P < 0.001, ****P < 0.0001, ns; not significant. Exact P values in Appendix Table S1.

To address this, we constructed strains expressing a PmrB variant with a partially deleted periplasmic domain (PmrB1 lacking amino acid residues 70-130) (Fig. 3B). It reported that the transmembrane domain is needed for antimicrobial peptide sensing in a different TCS sensor protein, PhoQ (Bader *et al*, 2005), we also constructed a strain expressing a PmrB variant with a partially deleted transmembrane domain 1 (PmrB2 lacking residues 23-29) (Fig. 3B). The PmrB1 strain exhibited significant survival defects upon exposure to H_2_O_2_ (Fig. 3C), suggesting that the periplasmic domain is required for H_2_O_2_ sensing. In contrast, bacterial survival was similar in strains harboring full-length PmrB or the truncated variant PmrB1 when treated with colistin (Fig. 3D). The survival of the PmrB2 strain was not affected by H_2_O_2_ treatment (Fig. 3C), but was significantly reduced when exposed to colistin (Fig. 3D). To test whether the PmrB variant localizes in the inner membrane of bacteria like the full-length PmrB protein, we performed subcellular localization analysis. It verified that PmrB1 and PmrB2 mainly localize to the inner membrane (IM) of *A. baumannii*, as indicated by the reduction of NADH (Appendix Fig. S5A-F) (Cian *et al*, 2020). Thus, the periplasmic domain of the PmrB sensor is required for oxidative stress sensing, and the transmembrane domain is required for colistin sensing (Fig. 3A).

Intriguingly, we identified four highly conserved histidine residues (His86, His89, His90, and His92) in the periplasmic domain of PmrB in *Acinetobacter* (Appendix Fig. S6A). These are specific to the *Acinetobacter* species, as determined by comparative amino acid alignment analysis (Appendix Fig. S6B). Notably, pathogenic *Acinetobacter* species, including *A. baumannii*, *A. calcoaceticus*, *A. lactucae*, *A. pittii*, and *A. haemolyticus*, possess these conserved four histidine residues (Appendix Fig. S6A). These species can cause pneumonia, urinary tract infections, and sepsis in humans (Cosgaya *et al*, 2019; Villalon *et al*, 2019), suggesting that conserved histidine residues may be critical for the virulence of pathogenic *Acinetobacter* species. Furthermore, we found that PmrB proteins from *Salmonella* or *Pseudomonas*, which lack this histidine box, could not restore H₂O₂ resistance in an *A. baumannii pmrB* mutant (Appendix Fig. S7B), even though these proteins can be well-expressed in an *A. baumannii pmrB* mutant strain by plasmid (Appendix Fig. S7A). Consistent with previous findings (Wosten *et al*., 2000), PmrA/PmrB is not involved in oxidative stress defense in *Salmonella*, whereas *oxyR* is involved in oxidative stress defense in this species (Appendix Fig. S7C). However, PmrB proteins from *Salmonella* or *Pseudomonas* restore resistance to antimicrobial peptides, such as colistin (Appendix Fig. S7D). Collectively, these observations led us to reason that histidine residues (His86, His89, His90, and His92) in the periplasmic domain of PmrB may be critical for oxidative stress sensing in *Acinetobacter*.

Multiple lines of evidence support the idea that histidine residues (His86, His89, His90, and His92) in the periplasmic domain of PmrB are critical for sensing oxidative stress. First, substitution of individual histidine residues or all four histidines with alanine decreased bacterial survival against H_2_O_2_ (Fig. 3E). Second, substitution of histidine residues did not affect colistin resistance (Fig. 3F). By contrast, the PmrB variant with substitution of a serine residue to alanine (PmrB*^Ser88^*) near the histidine residues could restore *A. baumannii* survival against H_2_O_2_. Third, the expression levels of both *hscA* and *ahpF1* were impaired in the *pmrB* mutant harboring PmrB with alanine-substituted four histidine residues upon H_2_O_2_ treatment compared to wild-type PmrB (Appendix Fig. S8A-B). In summary, four histidine residues (His86, His89, His90, and His92), referred to as the histidine box, are crucial for sensing oxidative stress but not for antimicrobial peptide (Fig. 3A).

### PmrB utilizes nickel as a cofactor with a histidine box for oxidative stress sensing

Next, it raised a question of how the histidine box in PmrB’s periplasmic domain senses oxidative stress. In many proteins, histidine residues participate in the binding of metal cations that play catalytic or structural functions (Dhakshnamoorthy *et al*, 2013). Also, the PmrB sensing domain has barrel structures, and they are often involved in oxidative stress sensing through metal cofactors in other enzymes (Perez-Dominguez *et al*., 2021; Tala *et al*., 2018). Therefore, we examined whether PmrB requires a metal cofactor for oxidative stress sensing with the histidine box.

Surprisingly, supplementation with nickel (Ni^2+^) in defined minimal medium without metal cations significantly enhanced the expression of both *hscB*-*hscA*-*fdx* and *ahpF1* in response to H_2_O_2_ (Fig. 4A-B), while other metals (Co²⁺, Mn²⁺, Zn²⁺) had no effect (Appendix Fig. S9A-C). Additionally, we detected Ni^2+^ using inductively coupled plasma mass spectrometry (ICP-MS), but not other metals, in the purified periplasmic domain (amino acid residues 30-140) of PmrB. By contrast, amounts of Ni^2+^ significantly reduced level in purified PmrB variant with the histidine box alanine substitution (PmrB*^His86,89,90,92^)* (Fig. 4C). Furthermore, a fluorescence-based dye for Ni^2+^ indicated that the amounts of Ni^2+^ were highly accumulated dependent on wild-type PmrB amounts (Fig. 4D). By contrast, Ni^2+^ detection was dramatically reduced in bovine serum albumin (BSA) and purified PmrB variant with the histidine box alanine substitution (PmrB*^His86,89,90,92^)* (Fig. 4D). Subsequently, adding Ni^2+^ (Fig. 4E), but not Co^2+^, Mn^2+^, or Zn^2+^ (Appendix Fig. S9D-F), increased resistance to oxidative stress in wild-type *A. baumannii*, consistent with Ni^2+^’s role in activating oxidative stress gene expression (Fig. 4A-B). By contrast, the *pmrB* null mutant remained highly sensitive to H_2_O_2_ regardless of Ni^2+^ (Fig. 4E).

**Figure 4.**
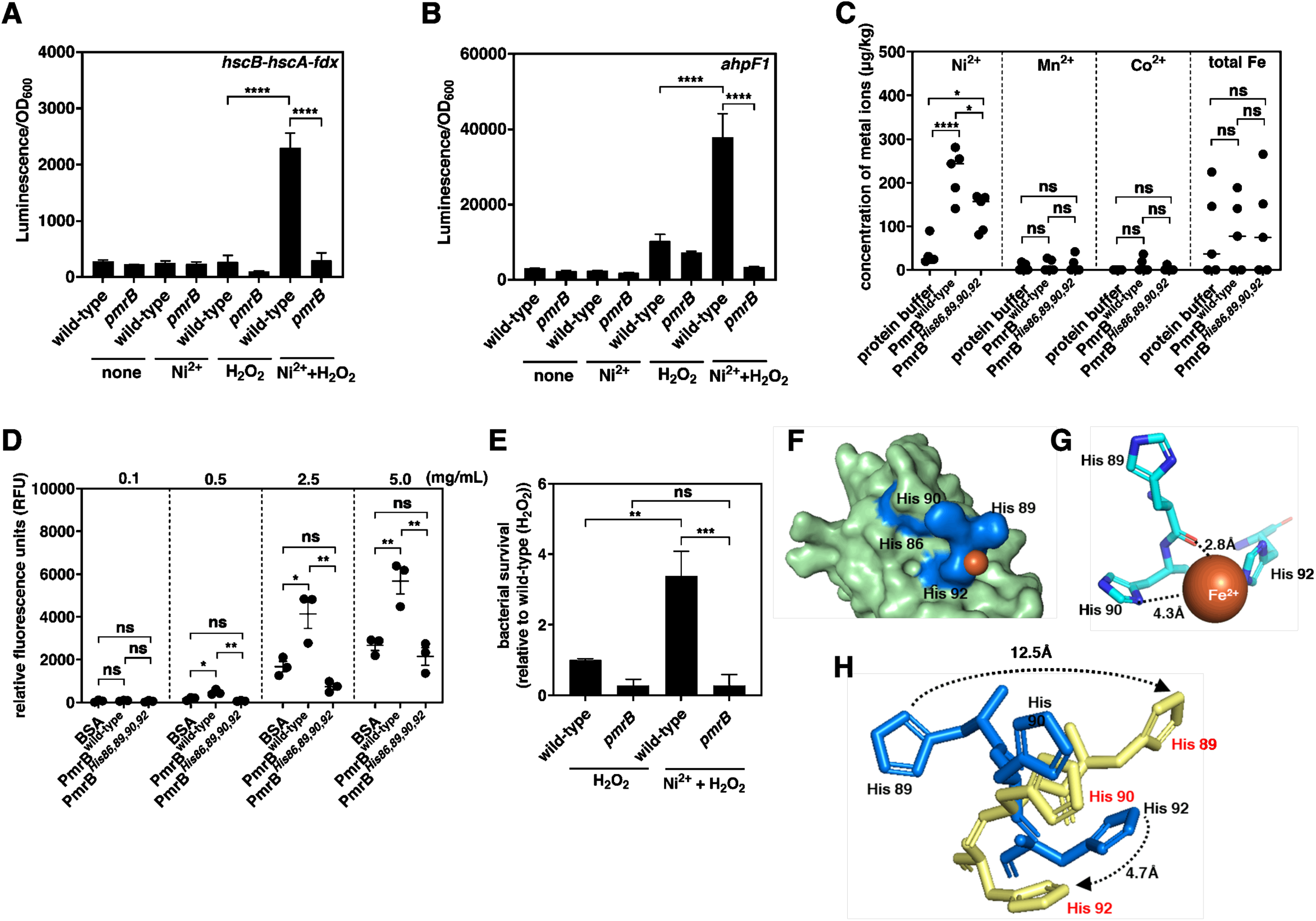
Nickel is a metal cofactor required for PmrB to sense oxidative stress. (A,B) Luminescence emission of *A. baumannii* 17978 carrying the luciferase reporter plasmid pLPV1Z fused with *hscB-hscA-fdx* promoter (A) or *ahpF1* promoter (B) at 2 h using M9 medium supplemented with 100 uM nickel (II) (Ni^2+^) and 10 mM H_2_O_2_. (C) Analysis of nickel (Ni^2+^), manganese (Mn^2+^), cobalt (Co^2+^), and total iron (total Fe) in wild-type PmrB protein fragment (PmrB^wild-type^) containing aa 30-140 and the same length PmrB fragment mutated in four histidine residues (PmrB*^His86,89,90,92^*) using inductively coupled plasma mass spectrometry (ICP-MS). (D) Quantification of nickel (Ni^2+^) in bovine serum albumin (BSA), wild-type PmrB protein fragment (PmrB^wild-type^) and PmrB fragment mutated in 4 histidine residues (PmrB*^His86,89,90,92^*) using the fluorescent dye Newport Green. In each protein samples, 4 concentrations (0.1, 0.5, 2.5, and 5.0 mg/mL) were used. (E) Relative survival of *A. baumannii* 17978 wild-type and *pmrB* after treatment with 10 mM H_2_O_2_ in the presence or absence of 100 uM nickel (Ni^2+^). (F-H) AlphaFold models of PmrB periplasmic domains with/without metal cofactor. (F) PmrB protein structure modelled by Alphafold3 with iron (Fe^2+^). The PmrB periplasmic domain (30-140) is depicted in green, the histidine box (His86, His89, His90 and His92) in blue and iron (Fe^2+^) in brown. (G) Distance between the histidine box (His89, His90 and His92) and iron (Fe^2+^) is measured by PyMol program. (H) Structural comparison between the histidine residues (His89, His90 and His92) with (blue color)/without (yellow color) iron (Fe^2+^). Distance between molecules is measured by PyMol program. Data are presented as means ± SD from n = 3 (A, B, D, and E) or n = 5 (C). *P* values were determined using two-way ANOVA with Tukey’s multiple comparision test (A, B, and E) or one-way ANOVA with Tukey’s multiple comparision test (C and D). *P < 0.05, **P < 0.01, ***P < 0.001, ****P < 0.0001, ns; not significant. Exact P values in Appendix Table S1.

Furthermore, we used AlphaFold3 to predict the PmrB structure with and without a metal cation. Since AlphaFold3 does not provide Ni²⁺ as a ligand, we substituted Fe²⁺, which has similar chemical features to Ni²⁺ (Abramson *et al*, 2024). Modeling showed that His89, His90, and His92 directly bind metal ions, while His86 may stabilize their interaction with nickel (Fig. 4F-G). Interestingly, PmrB structural conformation is changed by a metal cofactor (Fig. 4I). When a metal cofactor is present, His89 shifts rightward by ∼12.5 Å, and His92 moves downward by ∼4.7 Å (Fig. 4I, arrows from blue to yellow). This rearrangement enables His89, His90, and His92 to form a pocket for the Ni2+ cofactor, facilitating tight binding (Fig. 4I). Taken together, the Ni2+ cofactor with the histidine box is required for activation of gene expression for oxidative stress defense through PmrB in *A. baumannii*. Notably, this is the first report of a sensor protein in a two-component regulatory system using a Ni2+ cofactor with multiple histidine residues for oxidative stress sensing.

### Nickel oxidation drives redox-dependent structural reorganization of PmrB sensor

To detect oxidative stress molecules, the metal cofactor can be oxidized, triggering a conformational change in the sensor proteins that activates oxidative stress defense (Maroney & Ciurli, 2014; Pelmenschikov & Siegbahn, 2006). In *Bacillus*, PerR, a transcriptional regulator, contains metal-binding sites with multiple histidine residues (Lee & Helmann, 2006). The Fe^2+^ cofactor bound to histidine residues is oxidized by H_2_O_2_. This induces a conformational change in PerR, derepressing its target genes (Lee & Helmann, 2006). Also, we found that the PmrB structure reorganizes with a metal cofactor (Fig. 4I). Based on this, we hypothesized that oxidation of the metal cofactor by H_2_O_2_ in the histidine box may promote PmrB reorganization to enhance signal transduction. To test this hypothesis, we performed molecular dynamics (MD) simulations of apo-, Ni²⁺-and Ni³⁺-PmrB for 100 ns using bonded metal models (Bannwarth *et al*, 2019; Eastman *et al*, 2024; Maier *et al*, 2015; Wang *et al*, 2017).

Throughout all simulations, the coordination between nickel and the four histidine residues (His86, His89, His90, and His92) was stably maintained, confirming the integrity of the metal-binding geometry regardless of oxidation (Appendix Fig. S10A-B) (Maroney & Ciurli, 2014). Oxidation of the bound nickel from Ni²⁺ to Ni³⁺ drove a pronounced decreased structural stability in PmrB periplasmic domain (amino acid residues 29-136), with the Ni³⁺-PmrB reaching root mean square deviation (RMSD) values of ∼5.5 Å, and this is nearly 40% higher than either the apo- or Ni²⁺-PmrBs (∼4 Å) (Fig. 5A). Paradoxically, Ni³⁺-PmrB maintains tighter local geometry than apo-PmrB at the histidine box (Fig. 5B), indicating that oxidized nickel makes more unstable structure in overall PmrB protein, but Ni³⁺ is tightly bound to PmrB histidine box. Furthermore, per-residue root mean square fluctuation (RMSF) analysis revealed that Ni³⁺-PmrB displayed elevated flexibility across multiple regions, particularly around residues 68-85 and 108-136, while the binding site itself remained relatively ordered in all three conditions (Appendix Fig. S10C-D, yellow box) (Radivojac *et al*, 2007). Consistently, solvent accessible surface area (SASA) analysis indicated that the Ni³⁺-PmrB has higher SASA levels than other structures (Appendix Fig. S10E), which indicates that the Ni³⁺-PmrB structure is a more solvent-exposed conformation compared to Ni²⁺. Also, the radius of gyration analysis, which indicates a protein’s overall structural compactness and size, suggested that Ni²⁺-PmrB is the most compacted structure, while Ni³⁺-PmrB is the most expanded conformation (Appendix Fig. S10F) (McGibbon *et al*, 2015). Collectively, although oxidized nickel (Ni³⁺) strongly binds to the histidine box of PmrB, oxidation of the nickel cofactor promotes a change in protein conformation to the open form in PmrB.

**Figure 5.**
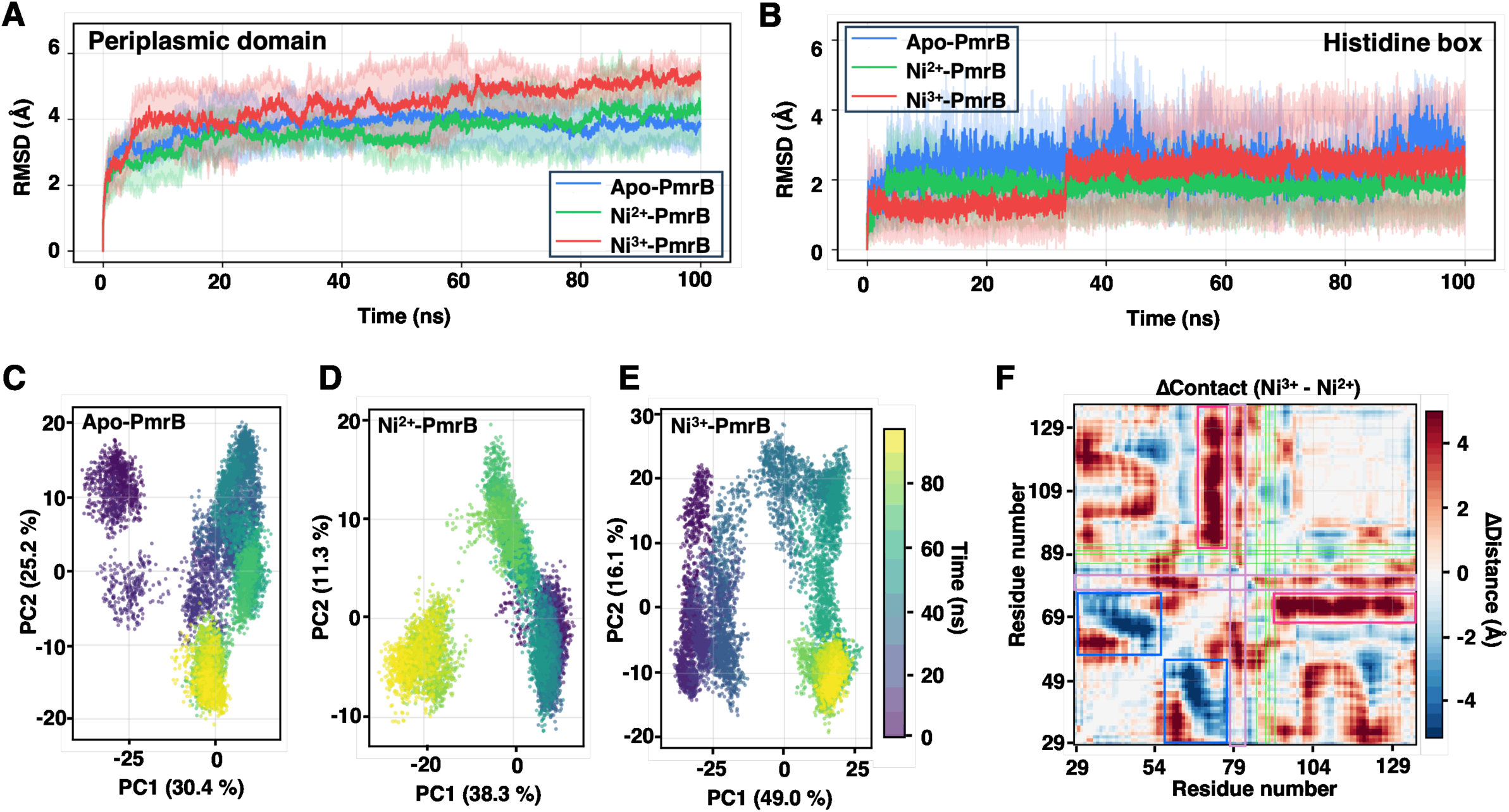
Oxidation of nickel cofactor promotes allosteric conformational change to active PmrA/PmrB system under oxidative stress conditions. (A, B) Root-mean-square deviation (RMSD) of the nickel-binding protein over 100 ns molecular dynamics (MD) simulations was performed to analyze to structural stability in PmrB periplasmic domain. (A) Overall Cα RMSD for the Apo (blue), Ni^2+^ (green), and Ni^3+^ (red) systems. (B) Binding site RMSD calculated for the four coordinating histidine residues (His86, His89, His90, and His92). Solid lines represent the mean values across three independent replicates (n = 3), and shaded regions indicate the standard deviation. The Ni^3+^-PmrB state exhibits the highest overall RMSD variability, while the binding site RMSD remains lowest in the Ni^3+^ system, suggesting tighter local coordination despite greater global structural fluctuations. (C-E) Directed conformational transitions of PmrB is measure by principal component analysis (PCA) of Cα atom trajectories. Projection onto the first two principal components (PC1 vs. PC2) for (C) Apo, (D) Ni^2+^, and (E) Ni^3+^ systems. Data points are colored by simulation time (purple: 0 ns; yellow: 100 ns). The percentage of variance explained by each component is indicated on the axes. The Apo state (PC1: 30.4%, PC2: 25.2%) exhibits broad conformational sampling, while the Ni^2+^ (PC1: 38.3%) and Ni^3+^ (PC1: 49.0%) states show progressively more directed conformational transitions along PC1. (F) Overall conformational change of PmrB is measure by differential contact map between Ni^3+^ and Ni^2+^ states. The heatmap displays the difference in average inter-residue Cα–Cα distances (ΔDistance = Ni^3+^ − Ni^2+^) averaged over three replicates using the last 50 ns. Red regions indicate increased inter-residue distances (expansion) and blue regions indicate decreased distances (compaction) in the Ni^3+^ state relative to Ni^2+^. Green lines demarcate the binding site residues (His86, His89, His90, and His92).

To investigate whether oxidized nickel promotes directional conformational change in PmrB, we performed principal component analysis (PCA) of the Cα trajectories (Amadei *et al*, 1993) using MDAnalysis (Michaud-Agrawal *et al*, 2011). It revealed that Ni³⁺ drives the protein toward a dramatically distinct conformational state (Fig. 5C-E). In the PCA plot, PC1 alone captured 49.0% of total structural variance in the Ni³⁺-PmrB (Fig. 5E), compared to just 38.3% for the Ni²⁺-PmrB (Fig. 5D) and 30.4% for apo-PmrB (Fig. 5C), respectively. Also, the initial structure (purple color) is clearly linked and directed to the final structure (yellow color) in Ni³⁺-PmrB (Fig. 5E). This indicates that nickel oxidation imposes a dominant, directional conformational transition on the PmrB protein (Fig. 5E). Subsequently, to determine which residues are reorganized by oxidized nickel in PmrB underlying this transition, we computed differential inter-residue contact maps from the last 50 ns of simulation using MD analysis (Michaud-Agrawal *et al*., 2011). The differential contact map (ΔContact=Ni³⁺-Ni²⁺) revealed a biphasic pattern of structural reorganization upon nickel oxidation (Fig. 5F). A prominent band of positive ΔDistance values (red) at residues ∼75–85 (purple box) indicated outward loop displacement in the Ni³⁺ state, consistent with elevated RMSF in this region (Appendix Fig. S10D). Additional long-range expansions appeared between residues ∼70-75 and ∼90–130 (Fig. 5F, pink box). By contrast, compensatory compaction (blue) between residues ∼30-50 and ∼55-78 (Fig. 5F, blue box) suggested a complex redistribution of inter-residue contacts, rather than simple global expansion, indicating an allosteric conformational change in PmrB upon nickel oxidation that might transfer the signal to the PmrA regulator. Intriguingly, the histidine box (green lines) resided at the boundary between expansion and compaction zones with near-zero ΔDistance values (Fig. 5F). This indicates that the local metal coordination geometry in the histidine box is preserved despite extensive surrounding reorganization. This is consistent with the low binding-site RMSD observed for Ni³⁺ (Fig. 5B). Taken together, single-electron oxidation of the nickel cofactor is sufficient to trigger a global conformational change of PmrB while preserving the integrity of the metal coordination shell. This is a structural hallmark of a redox-driven conformational switch mechanism (Maroney & Ciurli, 2014; Pelmenschikov & Siegbahn, 2006).

### PmrB forms response memory via sublethal oxidative stress that enables protection against lethal antimicrobial peptides and oxidative stress

In genetic regulatory systems, when an initial stimulus is removed, the bacterial cell can remain in the on state, a phenomenon known as response memory (Scanlon *et al*, 2025). Response memory, formed by an initial mild signal, primes the signal transduction system, dramatically enhancing the response to later signals (Lambert & Kussell, 2014). This mechanism is a critical adaptive strategy in environments that allows organisms to prepare for threats before lethal conditions occur.

During infection, pathogenic bacteria encounter sublethal oxidative stress from fluid flow in the bloodstream and airways (Forman *et al*., 2016; Padron *et al*., 2023), which are the initial sites of infection in sepsis and pneumonia, respectively. Later, bacteria face harsher stressors, such as high levels of oxidative stress and antimicrobial peptides released by innate immune cells, including macrophages and neutrophils (Herb & Schramm, 2021; Nguyen *et al*., 2017; Segal, 2005). In *A. baumannii*, the PmrA/PmrB system responds to multiple signals, including oxidative stress and antimicrobial peptides (Fig. 3). Notably, oxidative stress can enhance expression of both antimicrobial peptide defense genes, *pmrC* and *naxD* (Appendix Fig. S11), and oxidative stress defense genes (Fig. 2). By contrast, colistin, an antimicrobial peptide, did not induce oxidative stress defense genes (Fig. 2B and E, Appendix Fig. S4B and E). Given this, we hypothesized that sublethal oxidative stress at initial infection sites may signal the PmrA/PmrB system to prepare a second, stronger set of signals, such as lethal antimicrobial peptides or increased oxidative stress during infection, in *A. baumannii*.

To investigate this hypothesis, we exposed bacteria to sublethal concentrations of H_2_O_2_ (1, 5, or 10 µM) for 1 hour. Noted, bacteria could encounter these levels during infection (e.g., 1∼5 µM H_2_O_2_ in the bloodstream) (Forman *et al*., 2016; Padron *et al*., 2023). We then followed with high concentrations of H_2_O_2_ (30 mM) or colistin (3 µg/mL) for 2 hours (Fig. 6A). Remarkably, pretreatment with sublethal H_2_O_2_ dramatically increased expression of both oxidative stress defense genes (Fig. 6B and C; and Appendix Fig. S12A-B) and antimicrobial peptide resistance genes (Fig. 6D). This increase occurred in a PmrA-dependent manner. As expected, priming via sublethal ROS also significantly enhanced bacterial survival against high concentrations of both H_2_O_2_ (Fig. 6E) and colistin (Fig. 6F) compared to unprimed bacterial cells. By contrast, pretreatment with sublethal H_2_O_2_ doses in *oxyR* mutants did not enhance bacterial survival upon subsequent exposure to high H_2_O_2_ concentrations (Fig. 6G). This is consistent with a previous finding that inactivation of *oxyR* did not impair bacterial survival under oxidative stress (Appendix Fig. S3C) (Juttukonda *et al*., 2019). In addition, the histidine box of PmrB is essential for response memory function, because the priming effect from sublethal oxidative stress was completely absent in the histidine-substituted PmrB mutant (PmrB*^His86,89,90,92^*) in both oxidative and antimicrobial peptide resistance genes (Appendix Fig. S13). In sum, sublethal ROS forms a response memory of the PmrB sensor by priming that promotes responses to strong second signals, such as lethal oxidative stress or antimicrobial peptides.

**Figure 6.**
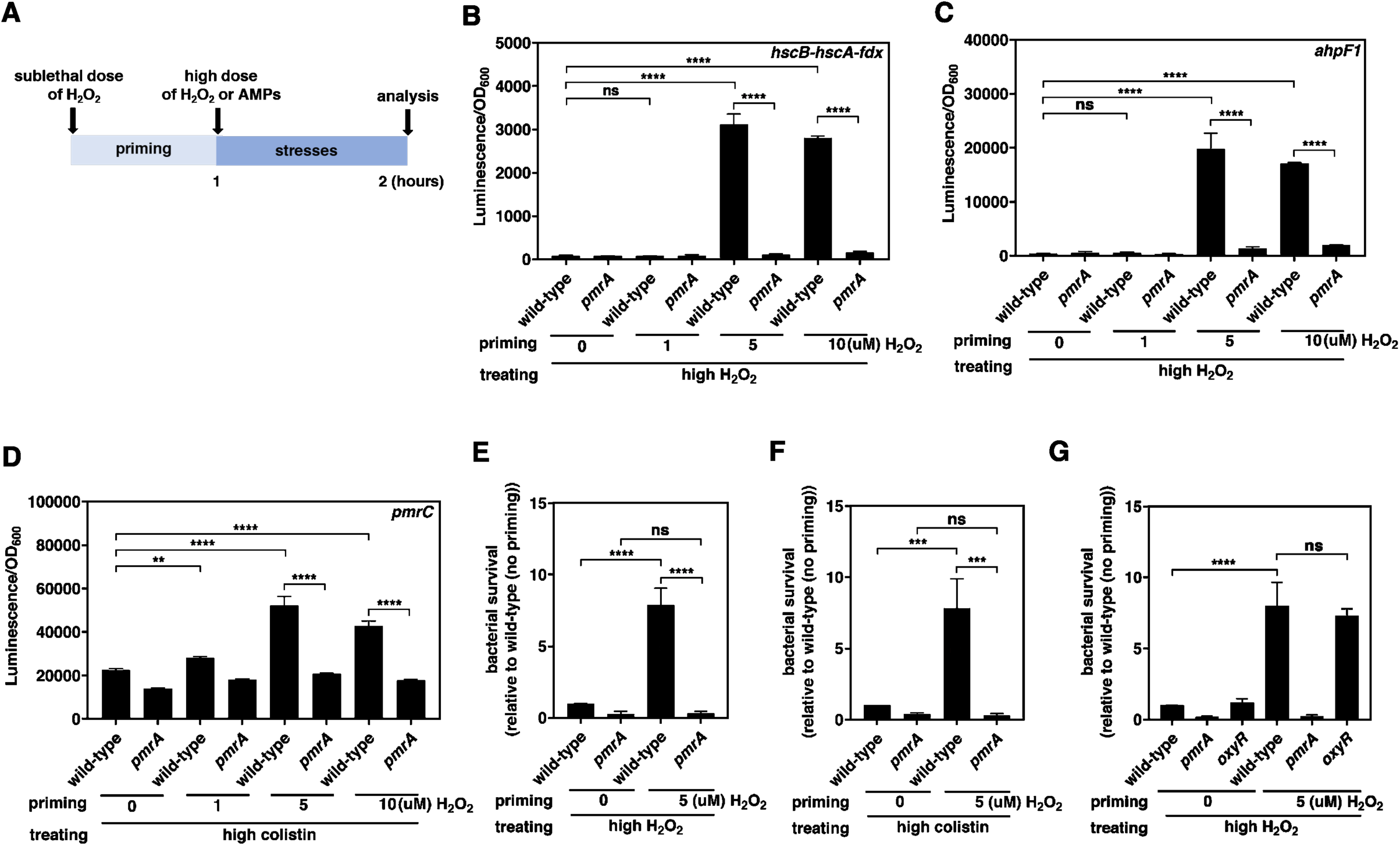
Priming effect via sublethal dose of H_2_O_2_ increased gene expression and bacterial survival under lethal ROS and antimicrobial peptide in *A. baumannii*. (A) Experimental design to test the priming effect of H_2_O_2_. Bacteria are treated with sublethal dose of H_2_O_2_ for 1 h and then treated with high dose of H_2_O_2_ or antimicrobial peptides (AMPs, colistin or LL-37) for 2 h. Gene expression and survival of bacteria were then measured. (B-D) Luminescence emission of *A. baumannii* 17978 wild-type and *pmrA* carrying the luciferase reporter plasmid pLPV1Z fused with *hscB-hscA-fdx* promoter (B), *ahpF1* promoter (C), or *pmrC* promoter (D) at 2 h in the presence 30 mM H_2_O_2_ (B,C) or 3 µg/mL colistin (D) after priming with different concentrations of H_2_O_2_(0, 1, 5, and 10 µM) for 1 h. (E, F) Relative survival of *A. baumannii* 17978 wild-type and *pmrA* after being primed with 5 µM H_2_O_2_ and treated with 30 mM H_2_O (E) or 3 µg/mL colistin (F). (G) Relative survival of *A. baumannii* 17978 wild-type and *oxyR* after being primed with 5 µM H_2_O_2_ and treated with 30 mM H_2_O. Data are presented as means ± SD from n = 4 (B, C, and D) or n = 3 (E, F and G). *P* values were determined using two-way ANOVA with Tukey’s multiple comparison test. **P < 0.01, ***P < 0.001, ****P < 0.0001, ns; not significant. Exact P values in Appendix Table S1.

Next, to test how long the response memory of PmrB is maintained by sublethal oxidative stress, we first exposed bacteria to sublethal 5 µM H_2_O₂ for 1 hour. Then, we removed H_2_O₂ and allowed the bacteria to rest for various durations. After resting, we administered high concentrations of H_2_O₂ (30 mM) or colistin (3 µg/mL) for 2 hours (Fig. 7A). We then measured expression of genes associated with oxidative stress defense and antimicrobial peptide resistance (Fig. 7A). Notably, the priming effect lasted for 30-90 minutes after removing the initial signal, sublethal H_2_O₂ (Fig. 7B-D). This may provide bacteria with a window of memory, enhancing cross-protection against strong stresses, such as lethal oxidative stress and antimicrobial peptides, during infection. To rule out that the loss of response memory effect results from bacterial growth, we counted colony forming units (CFUs) before and after removing the sublethal initial signal. As expected, CFU count did not significantly change (Appendix Fig. S14). This shows that cellular component dilution via cell division is not the reason for the loss of response memory after initial signal removal. Taken together, the PmrA/PmrB system’s response memory to sublethal ROS provides a window to prepare defenses against both lethal oxidative stress and antimicrobial peptides in *A. baumannii*.

**Figure 7.**
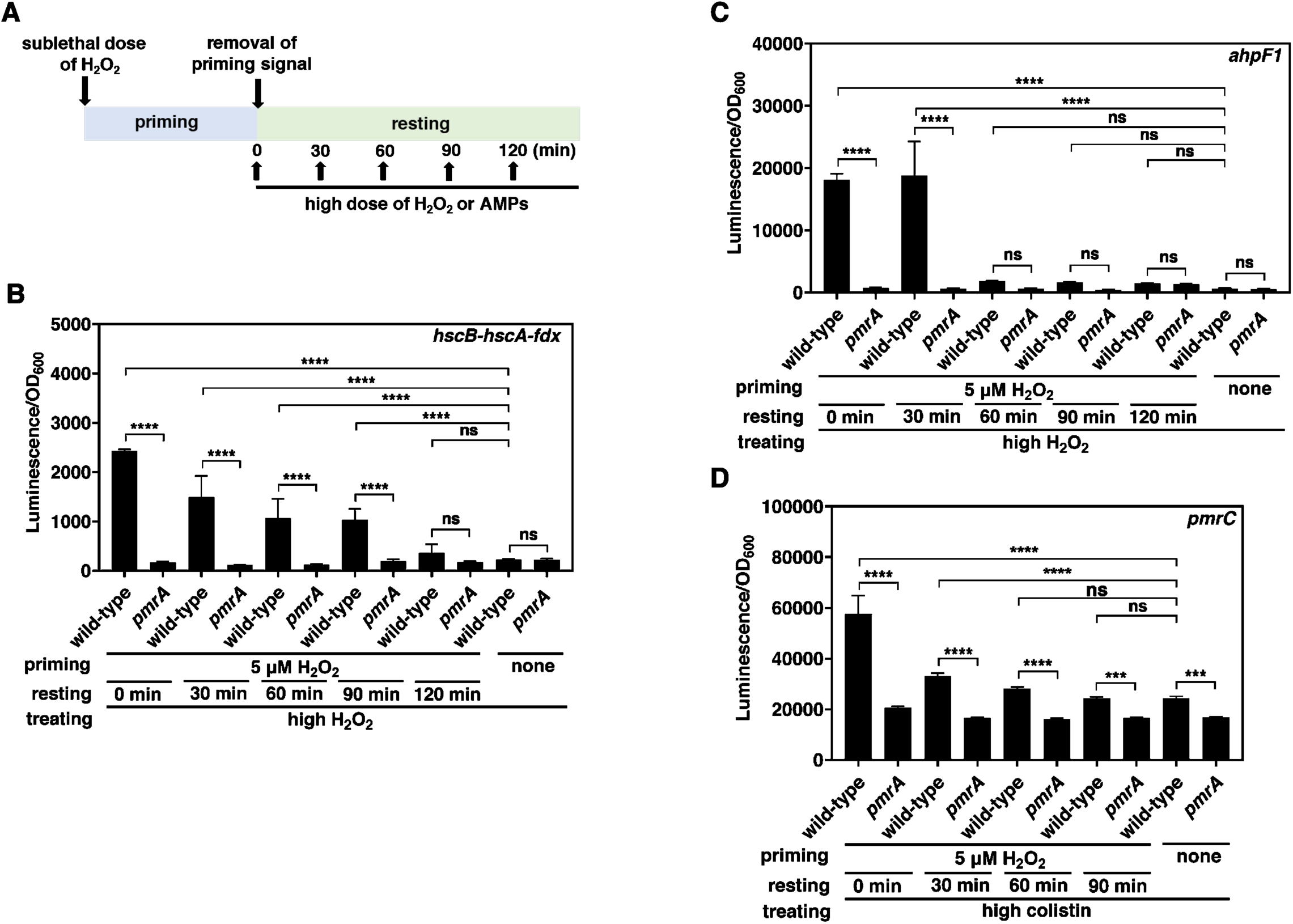
PmrB maintains the priming effect for a certain time after the removal of the initial signal that confers response memory. (A) Experimental design to test how long the priming effect of H_2_O_2_ lasts. Bacteria are treated with sublethal dose of H_2_O_2_ for 1 h, rested for different times after the priming signal is removed, then treated with high dose of H_2_O_2_ or antimicrobial peptide colistin for 2 h. Gene expressions were then measured. (B-D) *A. baumannii* 17978 wild-type and *pmrA* carrying the luciferase reporter plasmid pLPV1Z fused with *hscB-hscA-fdx* promoter (B), *ahpF1* promoter (C), or *pmrC* promoter (D) were primed with 5 µM H_2_O_2_ for 1 h. The primed bacteria were washed with warm LB broth to remove the prime signal and rested for different times (0, 30, 60, 90, and 120 min) before being treated with high doses of H_2_O_2_ (30 mM) (B and C) or colistin (3 µg/mL) (D). Luminescence emission was measured 2 h post-treatment with H_2_O_2_ (B and C) or colistin (D). Data are presented as means ± SD from n = 4. *P* values were determined using two-way ANOVA with Tukey’s multiple comparision test (B, C, and D). ***P < 0.001, ****P < 0.0001, ns; not significant. Exact P values in Appendix Table S1.

### Oxidative stress sensing of PmrB via the histidine box is necessary for virulence in the host

To examine whether oxidative stress sensing by PmrB is critical for virulence in the host, we intratracheally infected mice with wild-type and *pmrB* mutant *A. baumannii* (Fig. 8A). The wild-type strain induced severe lung inflammation, while the *pmrB* mutant caused only mild inflammation (Fig. 8B and C). As neutrophils are the first responding cells to pathogen-induced tissue inflammation (Davey *et al*, 2011), we investigated whether PmrB-mediated *A. baumannii* infection can modulate neutrophil infiltration (Appendix Fig. S15A). We observed significantly fewer neutrophils in the lungs of mice infected with the *pmrB* mutant compared to wild-type-infected mice using *Ly6g*^cre^:R26^LSL-TdTomato^ reporter mice (Appendix Fig. S15B), which facilitates the identification of neutrophils by fluorescent microscopy (Hasenberg *et al*, 2015) (Fig. 8D and E). Also, the number of neutrophils was significantly higher in mice infected with *pmrB* mutant harboring plasmid with wild-type PmrB compared to the mutant with empty vector (Fig. 8F). Moreover, we determined that the histidine box for oxidative stress sensing is critical for lung inflammation and neutrophil infiltration, because bacteria with PmrB containing histidine-to-alanine substitutions (PmrB*^His86,89,90,92^*) showed only partial neutrophil recruitment (Fig. 8F). Furthermore, bacterial counts in lung tissues were significantly reduced in mice infected with *pmrB* mutant compared to wild-type (Fig. 8G). In addition, bacterial counts are restored by expression of wild-type PmrB from a heterologous promoter in the *pmrB* mutant strain. By contrast, bacteria with PmrB containing histidine-to-alanine substitutions (PmrB*^His86,89,90,92^*) did not restore bacterial counts in lung tissue (Fig. 8G).

**Figure 8.**
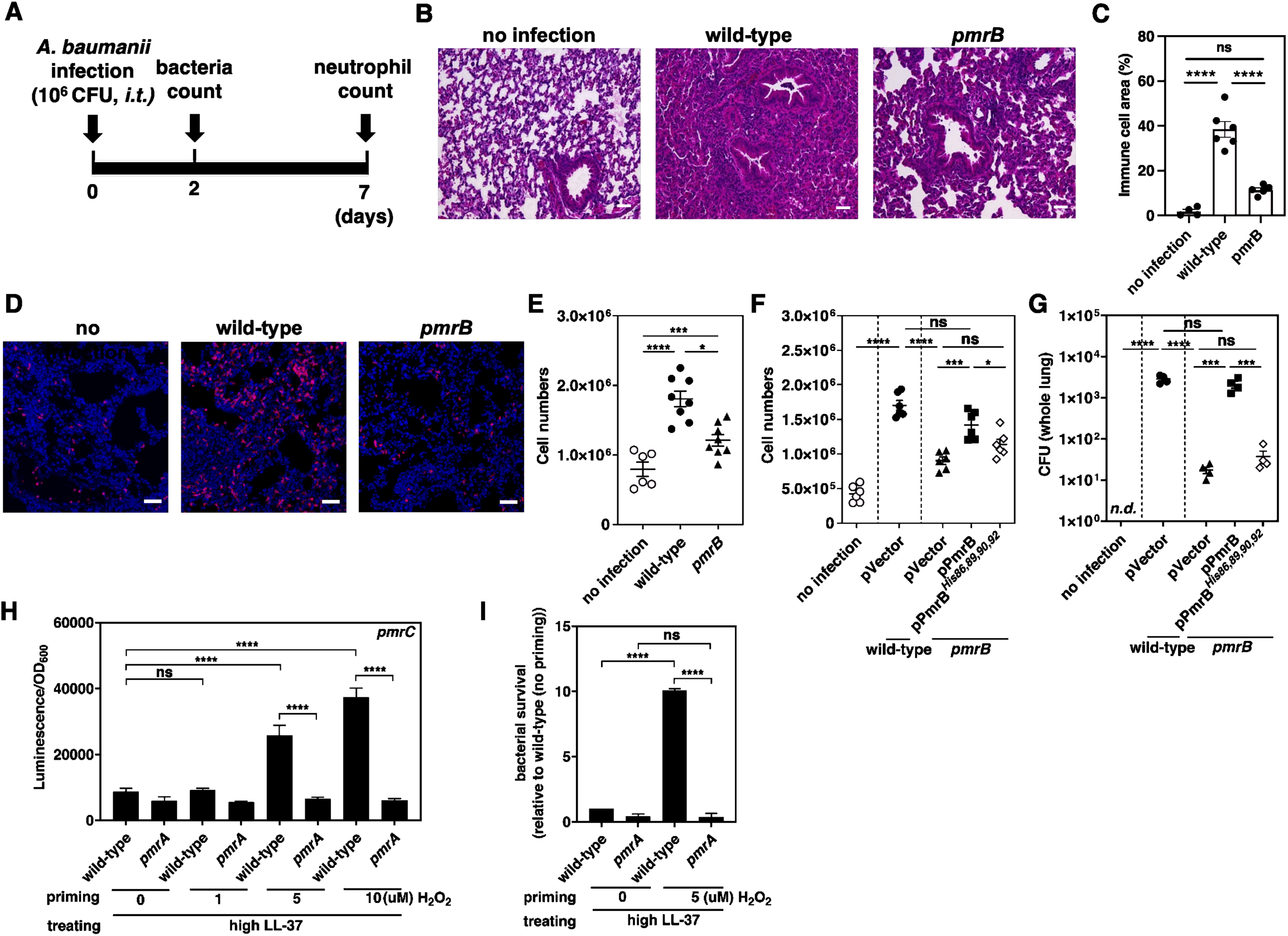
Histidine residues on the periplasmic domain of PmrB is crucial for the survival of *A. baumannii* inside immune cells and murine model. (A) Scheme for experimental design using a mouse model. 10^6^ CFU of *A. baumannii* strains intratracheally infected with mice, then bacterial colony forming unit (CFU) will be measured after 2 days post infection, and tissue inflammation of infected mice will be assessed by histology, immunohistochemistry and flowcytometric analysis after 7 days post infection. (B) H&E staining images of lung sections from naïve mice, wild-type *A. baumannii*-infected mice, and *pmrB*-infected mice. N = 4-6 mice per group; representative image from two independent experiments, scale bar; 50 µm). (C) Quantification of immune cell areas in H&E-stained images shown in (B). The area was quantified in two different regions of H&E-stained images per mouse (*n* = 5 per group; two-way ANOVA with Tukey’s multiple comparison test). (D) Immunohistochemimal analysis of lung sections from naïve Ly6gcre:R26LSL-TdTomato mice, wild-type or *pmrB*-infected Ly6gcre:R26LSL-TdTomato mice. (n=3-5 mice per group; data are representative images from two independent experiments). (E) Quantification of neutrophils in the lungs of non-infected mice, wild-type or *pmrB*-infected mice (n=6-8 mice per group; data are pooled from two independent experiments). (F) Flow cytometric quantification of neutrophils in the lungs of non-infected mice, or mice infected with wild-type harboring the plasmid vector (pWH1266) and *pmrB* harboring the plasmid vector, the pmrB-expressing plasmid (pPmrB), or the plasmid expressing *pmrB* mutated in histidine residues (pPmrB*^His86^*, pPmrB*^His89^*, pPmrB*^His90^*; pPmrB*^His92^* or pPmrB*^His86,89,90,92^*) (n=6 per group; data are pooled from two independent experiments). (G) Bacterial count in the lungs of non-infected mice, or mice infected with wild-type harboring the plasmid vector (pWH1266) and *pmrB* harboring the plasmid vector, the pmrB-expressing plasmid (pPmrB), or the plasmid expressing *pmrB* mutated in histidine residues (pPmrB*^His86^*, pPmrB*^His89^*, pPmrB*^His90^*; pPmrB*^His92^*or pPmrB*^His86,89,90,92^*) (n=4 per group; data are pooled from two independent experiments; n.d.; not detected). (H) Luminescence emission of *A. baumannii* 17978 wild-type and *pmrA* carrying the luciferase reporter plasmid pLPV1Z fused with *pmrC* promoter at 2 h in the presence of 10 µg/mL LL-37 after priming with different concentrations of H_2_O_2_ (0, 1, 5, and 10 µM) for 1 h. Data are presented as means ± SD from n = 4. (I) Relative survival of *A. baumannii* 17978 wild-type and *pmrA* after being primed and treated with 10 µg/mL LL-37. Data are presented as means ± SD from n = 3. *P* values were determined using one-way ANOVA with Tukey’s multiple comparision test (C, E, F, and G), or two-way ANOVA with Tukey’s multiple comparision test (H and I). * P<0.05, ** P <0.01, *** P<0.001, **** P<0.0001, ns; not significant. Exact P values in Appendix Table S1.

After neutrophil recruitment in response to bacterial infection, neutrophils can kill bacteria by secreting specific antimicrobial peptides, such as LL-37 (Neumann *et al*, 2014). In agreement with this, it raised the question of whether PmrB’s response memory to sublethal ROS is critical for bacterial protection from neutrophil-secreted antimicrobial peptides. Exposure to sublethal H_2_O_2_ significantly increased expression of antimicrobial peptide resistance gene *pmrC* under LL-37 treatment (Fig. 8H). By contrast, inactivation of *pmrA* did not prime *pmrC* expression in response to sublethal ROS under LL-37 treatment (Fig. 8H). In addition, priming by sublethal ROS dramatically increased bacterial survival to LL-37, in a PmrA-dependent manner (Fig. 8I).

Next, to test whether ROS sensing by PmrB in *A. baumannii* is critical for survival against innate immune cells such as macrophages and neutrophils, we measured bacterial survival within macrophage-like cells after infection. Survival rate of wild-type *A. baumannii at* 6 h post-phagocytosis was higher than that of *pmrA* or *pmrB* mutant strains inside macrophages (Appendix Fig. S16A). Moreover, bacterial survival was significantly higher in a *pmrA* or *pmrB* mutant harboring a plasmid with their respective *pmrA* or *pmrB* gene expressed from a heterologous promoter compared to an isogenic strain with the empty vector (Appendix Fig. S16B). Lastly, a strain expressing a PmrB sensor with the four histidine residues substituted to alanine (PmrB*^His86,89,90,92^*) failed to restore survival to a *pmrB* mutant (Appendix Fig. S16B). By contrast, the PmrB variant with substitution of a serine residue to alanine (PmrB*^Ser88^*) near the histidine residues could restore *A. baumannii* survival inside macrophages (Fig. S16B). Together, these results demonstrate that oxidative stress sensing and PmrA/PmrB response memory are crucial for *A. baumannii* virulence and survival in the host during infection.

### Oxidative stress sensing via the histidine box in PmrB is determinant for hypervirulence in clinically isolated carbapenem-resistant *A. baumannii*

The World Health Organization (WHO) designates carbapenem-resistant *A. baumannii* (CRAB) as a top-priority pathogen due to its critical health threat by multidrug resistance. Previously, we reported that hypervirulent CRAB, especially those isolated from bacteremia patients in the intensive care units of the hospital, cause death within 3 days of the initial positive blood culture, which is defined as early mortality (Lee *et al*, 2024). We determined here that response memory and oxidative stress sensing by the histidine box of PmrB are essential for *A. baumannii* virulence in the infected host (Fig. 7 and Fig. S16). Thus, we hypothesized that the histidine box in PmrB may be a hypervirulence determinant in clinically isolated CRAB strains.

To test this hypothesis, we used the laboratory strain ATCC 17978, which was used in all of the studies above. Also, a hypervirulent clinically isolated CRAB strain A0062, which caused early mortality, and a low-virulence clinically isolated *A. baumannii* strain A0075, which did not cause patient death, were used (Lee *et al*., 2024). Interestingly, the histidine box is conserved in the hypervirulent clinically isolated CRAB strain A0062 and the laboratory strain ATCC 17978, but absent in the low-virulence clinically isolated A0075 strain (Fig. 9A). Second, treatment of the low virulent clinical isolate *A. baumannii*A0075 strain with 30 mM H_2_O₂ completely abolished its survival (Fig. 9B). This agrees with the essential role of the histidine box in oxidative stress sensing and defense. In contrast, the laboratory strain ATCC 17978 could survive exposure to 30 or 60 mM H_2_O₂ (Fig. 9B and C). Noted, the hypervirulent CRAB A0062 strain is more resistant to oxidative stress than the laboratory strain, because a hypervirulent clinical isolate, CRAB A0062 strain, could resist very high concentrations of H_2_O₂ (up to 120 mM) (Fig. 9B-E). Thus, the hypervirulent clinically isolated CRAB A0062 strain, may have additional mechanisms to defend against high oxidative stress. Collectively, the histidine box of PmrB is essential for defense against oxidative stress in clinically isolated *A. baumannii*.

**Figure 9.**
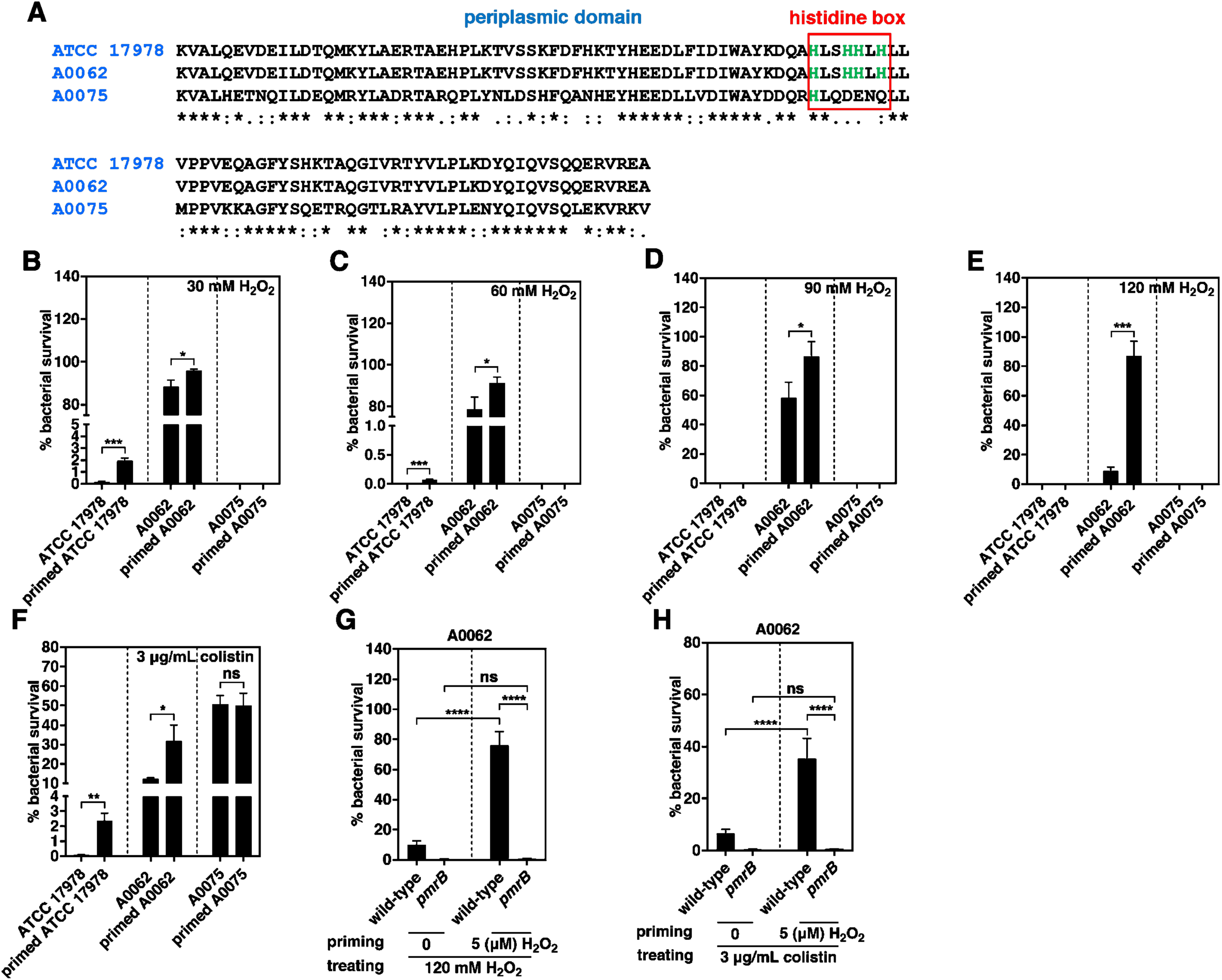
Response memory via PmrB-mediated H_2_O_2_ sensing is the virulent determinant of *A. baumannii* clinical isolates. (A) Amino acid alignment of PmrB periplasmic domains of laboratory strain *A. baumannii* ATCC 17978, high virulent CRAB (A0062), and low virulent *A. baumannii* (A0075). (B-F) Percentage of survival of *A. baumannii* ATCC 17978, A0062, and A0075 after being primed with or without 5 µM H_2_O_2_ for 1 h and treated with high concentrations of H_2_O_2_ (B-E) or colistin (F) for 2 h. (G and H) Percentage of survival of high virulent CRAB A0062 wild-type and *pmrB* mutant after being primed and treated with high concentrations of H_2_O_2_ (G) and colistin (H). Data are presented as means ± SD from n = 3. *P* values were determined using unpaired Student’s t-test (B-F) or two-way ANOVA with Tukey’s multiple comparison test (G and H). *P < 0.05, **P < 0.01, ***P < 0.001, ****P < 0.0001, ns; not significant. Exact P values in Appendix Table S1.

Furthermore, response memory of PmrB via a sublethal 5 µM H_2_O₂ in the hypervirulent clinically isolated CRAB A0062 significantly enhanced bacterial survival when exposed to high concentrations of H_2_O₂ and colistin later (Fig. 9B-F). In addition, inactivation of *pmrB* in the hypervirulent clinically isolated CRAB A0062 significantly reduced bacterial survival by removing the priming effect mediated by sublethal ROS (Fig. 9G and H). Taken together, our results strongly indicate that response memory via the histidine box of PmrB is an indispensable virulence determinant in the clinically isolated CRAB strains.

## Discussion

We have established that oxidative stress sensing by the PmrA/PmrB regulatory system is critical for full activation of *A. baumannii* virulence (Fig. 10). Importantly, exposure to sublethal concentrations of oxidative stress produces response memory and dramatically enhances resistance against both lethal oxidative stress and antimicrobial peptides by the PmrA/PmrB system, revealing a key mechanism that enables full bacterial virulence in *A. baumannii*. Pathogenic bacteria continuously encounter sublethal reactive oxygen species (ROS) from fluid flow in the bloodstream and airways during infection (Forman *et al*., 2016; Padron *et al*., 2023). In agreement with this, response memory to sublethal ROS concentrations is critical for understanding how bacterial pathogens prepare for lethal stresses later by fine-tuning their signal transduction systems (Fig. 10). Notably, we identified four histidine residues and a nickel cofactor (Ni^2+^) that sense oxidative stress molecules. Oxidation of nickel (Ni^3+^) by ROS molecules promotes an allosteric conformational change that activates the signal transduction system. Furthermore, we have determined that this mechanism is also important to hypervirulence in clinically isolated carbapenem-resistant *A. baumannii*, which is the most critical pathogen designated by the WHO. Therefore, we first report that PmrB forms a sensor of sublethal oxidative stress via a nickel cofactor and the histidine box, thereby conferring full virulence during infection.

**Figure 10.**
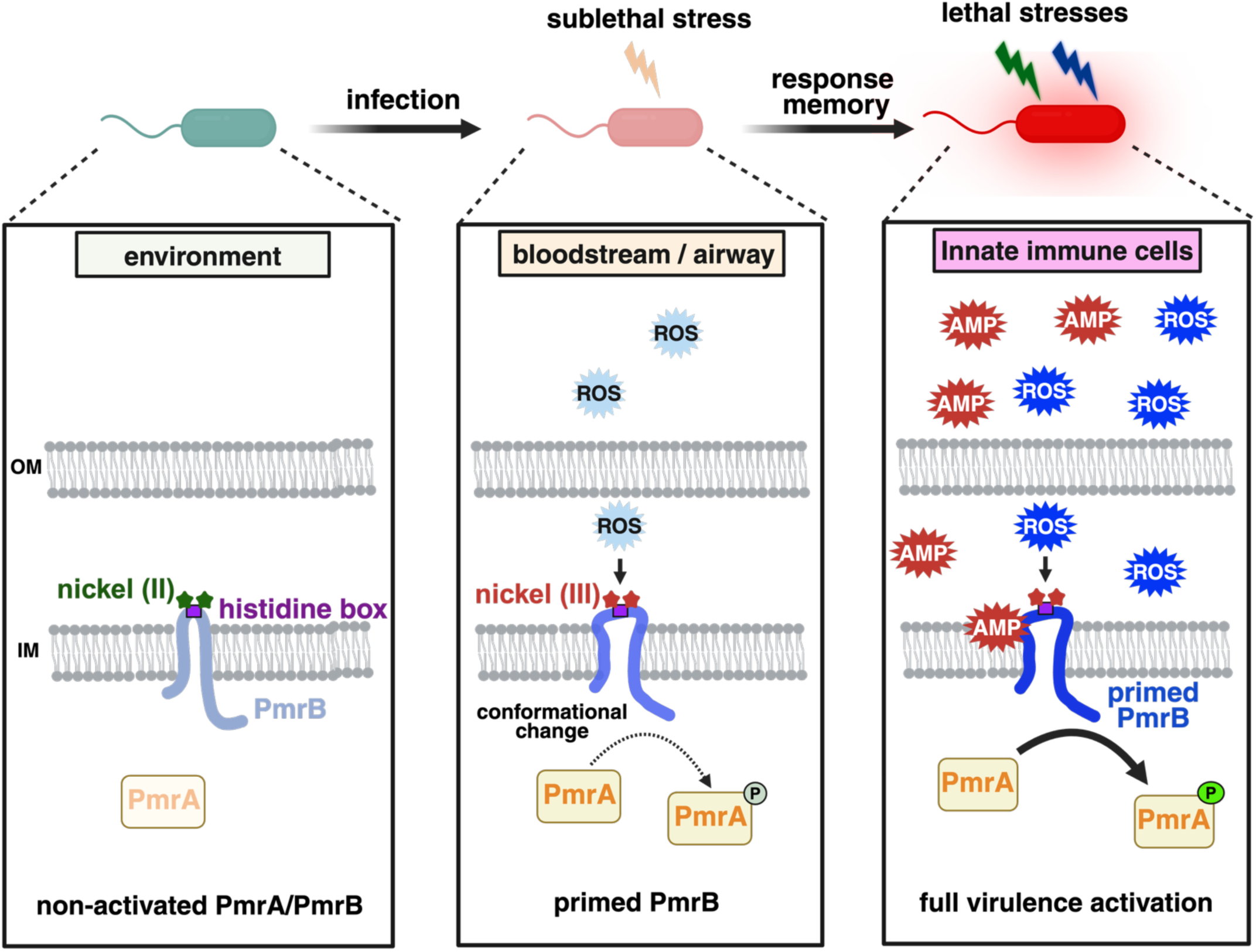
Sublethal ROS primes the PmrA/PmrB system that confers full activation of virulence by resistance to lethal stresses in *A. baumannii*. (Left) The PmrA/PmrB contains nickel (Ni^2+^) as a metal cofactor in the histidine box. Under normal conditions, the PmrA/PmrB system turns off because it does not sense and respond to stress. (Middle) During infection, bacteria usually encounter sublethal concentrations (1–10 µM) of reactive oxygen species (ROS such as H_2_O_2_) in the bloodstream or airways. Sublethal H_2_O_2_ acts as an initial signal and is detected in the periplasm by the sensor kinase PmrB, which relies on its histidine box and nickel oxidations (Ni^3+^). Once this signal is received, PmrB forms response memory, preparing the cell for upcoming harsh conditions. (Right) After bacterial infection, innate immune cells are recruited to kill bacteria. In innate immune cells, when exposed to lethal H_2_O_2_ or antimicrobial peptides (AMP such as colistin and LL-37), primed PmrB detects these signals and strongly up-regulates the defense system via PmrA, much more than in non-primed cells. As a result, bacteria gain increased resistance to lethal oxidative stress and antimicrobial peptides. This mechanism ultimately promotes hypervirulence of *A. baumannii* within the host.

Sensor proteins in TCS utilize specific amino acid residues in their sensory domains but typically do not require cofactors. Surprisingly, the PmrB sensor employs a nickel cofactor, enabling bacteria to sense and respond to oxidative stress. Metal cation cofactors are required for oxidative stress sensing in all three domains of life (Costa & Moradas-Ferreira, 2001; Lee & Helmann, 2006; Pedone *et al*, 2004). Specifically, nickel is a ubiquitous metal found in soil, water, and the human body (Angon *et al*, 2024). Nickel is essential for the survival and pathogenesis of pathogenic bacteria during infections (Zambelli *et al*, 2016). Indeed, nickel can be involved in ROS defense as a cofactor, such as in superoxide dismutase (SOD), a key enzyme in oxidative stress defense (Barondeau *et al*, 2004). Notably, we demonstrated that nickel could bind to a histidine-rich motif on the sensor domain of PmrB to sense oxidative stress (Fig. 4). In addition, using simulation analysis of molecular dynamics (MD), oxidative stress can oxidize nickel cofactor, then oxidized nickel (Ni^3+^) promotes to conformational change of PmrB periplasmic domain to active signal transduction system (Fig. 5). Furthermore, we determined which residues are involved in the conformational change of the PmrB protein by nickel cofactor oxidation (Fig. 5F). Therefore, the nickel cofactor bound to histidine residues forms a key pocket in PmrB to sense and respond to oxidative stress in *A. baumannii*, because allosteric effects via conformational change could be a hallmark of redox-driven sensing mechanisms in proteins (Maroney & Ciurli, 2014; Pelmenschikov & Siegbahn, 2006).

The priming effect via response memory of the signal transduction system, in which exposure to an initial signal enhances the response to subsequent signals, has been well documented in mammalian and plant immune cells, such as IFN-γ priming of innate immune cells (Borges da Silva *et al*, 2015; Wu *et al*, 2014; Hilker & Schmulling, 2019). Our findings reveal that pathogenic bacteria have evolved a parallel strategy through their signal transduction systems to survive and proliferate in the host (Fig. 10). Sublethal micromolar concentrations of ROS act as initial signals for *A. baumannii* (Fig. 6). These concentrations are naturally encountered in host tissues such as the bloodstream and airways (Forman *et al*., 2016; Padron *et al*., 2023). The initial signals heighten the sensitivity of the PmrA/PmrB system (Fig. 6). Remarkably, our data demonstrate that signal priming via response memory generates a survival advantage against both lethal oxidative bursts and antimicrobial peptides (Fig. 6). This effect persists for 30-90 minutes (Fig. 7), providing sufficient time for bacteria to adapt during critical phases of infection (Fig. 8). In pathogenic bacteria, it is poorly understood how the signal transduction system’s response memory prepares it to withstand the extreme stresses encountered during infection or phagocytosis by innate immune cells. In addition, recently, it reviewed the concept of bacterial memory and suggested that there is a gap in the key molecular mechanisms, even though evidence of bacterial memory continues to emerge (Scanlon *et al*., 2025). In this study, we provide a molecular mechanism of response memory in bacteria using an analytical chemical tool, biochemical analysis and machine learning based *in slico* simulation.

Currently, carbapenem-resistant *A. baumannii* (CRAB) is one of the most dangerous pathogens worldwide (Amaya-Villar & Garnacho-Montero, 2019; Mancuso *et al*, 2021). In addition, CRAB is a multidrug-resistant bacterium, and there are very limited antibiotic options to eliminate it in a clinical setting. Antimicrobial peptide drugs such as colistin are the last-line antibiotics against multidrug-resistant CRAB, but resistance is emerging (Falagas & Kasiakou, 2005; Novovic & Jovcic, 2023). Here, we discovered that a response memory of TCSs is essential for the full activation of the defense system against multiple strong stressors, including the last-line antibiotics colistin against CRAB in *A. baumannii* during infections (Fig. 10). Furthermore, four histidine residues and a nickel cofactor are determinants for hypervirulent clinically isolated *A. baumannii* (Fig. 9). Thus, disrupting the nickel-dependent sensing of the PmrB system could potentially compromise bacterial virulence and restore susceptibility to existing antibiotics, presenting a new approach to combat multidrug-resistant CRAB infections.

## Materials and Methods

### Bacterial strains and growth conditions

Bacterial strains and plasmids employed in this study are listed in Appendix Table S2. Unless otherwise indicated, experiments were performed with *A. baumannii* strain ATCC 17978 and its isogenic mutants. For routine cultivation and genetic manipulation, *A. baumannii*, *E. coli* and *Salmonella* were grown at 37°C in Luria Bertani (LB) broth or on LB agar (BD Difco). For reporter assays with metal ions, M9 medium (15 g/liter Na_2_HPO_4_, 1.5 g/liter NaCl, 7.5 g/liter KH_2_PO_4_, 5 g/liter NH_4_Cl, 0.4% glucose, 0.4% casamino acid, 0.1 mM CaCl_2_, and 2 mM MgSO_4_) was used. For selection of recombinants or maintenance of plasmids in *A. baumannii*, *E. coli* and *Salmonella*, antibiotics were supplied at the following concentrations: ampicillin, 50µ g/mL; apramycin, 100 µ g/mL; 100 carbenicillin, 100 µ g/mL; chloramphenicol, 25 µg/mL; gentamicin 10 µg/mL; and kanamycin, 50 µg/mL.

### Cell lines

The J774A.1 mouse macrophage cell line was cultured in Dulbecco’s Modified Eagle Medium (DMEM) High Glucose (Welgene) supplemented with 10% heat-inactivated fetal bovine serum (Welgene) at 37°C and 5% CO_2_.

### Mouse models

All animal studies were approved from the Institutional Animal Care and Use Committee (IACUC) at the University of Illinois at Chicago (UIC). Wild-type (WT) mice (C57B6/J, Jax:00664) and R26LSL-Tomato mice (Jax: 007909) were obtained from The Jackson Laboratory and Ly6gCre Catch-up mice were generously provided by Dr. Matthias Gunzer. Catch-up mice were crossed with R26LSL-Tomato mice to specifically label neutrophils.

### Construction of chromosomal mutants and plasmids in A. baumannii

Primers used in this study are listed in Appendix Table S3. Chromosomal mutants (HN9, HN13, HN17, HN160, HN164, HN208, HN330, HN502, and HN503) and wild-type with PmrB-3xFLAG (HN19) were generated using standard protocols described previously (Tucker *et al*, 2014). Briefly, the FRT-flanked kanamycin resistance determinant of pKD4 was amplified using primers bearing ∼50 bp of homology to the region flanking the gene of interest. This PCR product was electroporated into competent *A. baumannii* carrying pAT02 plasmid, which expresses the RecAb recombinase. Mutants were selected on LB agar plates with kanamycin, and integration of the resistance marker was confirmed by PCR. To remove kanamycin resistance cassette, electrocompetent mutants were transformed with pAT03 plasmid, which expresses the FLP recombinase. All mutant strains were confirmed by PCR and sequencing.

Complementation of *pmrA* or *pmrB* mutants (except Appendix Fig. S14A, B, and C) was performed *in trans* using the vector pWH1266 (Hunger *et al*, 1990). The gene and its native promoter were amplified using genomic DNA from wild-type ATCC 17978 as template and the primers listed in Appendix Table S3. pWH1266 was digested with BamHI and SalI (New England Biolabs). The purified PCR product and cut plasmid were joined using NEB Builder Hifi Assembly (New England Biolabs) at 50°C for 15 min. The resulting assembly was transformed into chemically compenent DH5α by heat shock, and successful transformants were selected on LB agar plates with ampicillin following overnight growth at 37°C. The constructs were confirmed by PCR and sequencing prior to introduction of this vector, as well as the native empty vector, to the mutants. Successful transformants were selected on LB agar plates with ampicillin. Complementation of *pmrB* mutant in Appendix Fig. S14A, B, and C was performed using the vector pJMP3653 (Bacon *et al*, 2024). The same procedure was used, except that pJMP3653 was cut with NcoI and BamHI and successful transformants were selected on LB agar plates with kanamycin. 25 µM IPTG was used to induce expression of *pmrB*.

To construct *pmrB* mutant harboring pPmrB-3xFLAG, pPmrB1-3xFAG, or pPmrB2-3xFLAG (HN283, HN287, HN505 respectively), the same procedure for making complementation has been done except that genomic DNA from wild-type with PmrB-3xFLAG (HN19) was used as template.

### Construction of insertional mutants in A. baumannii clinical isolates

*pmrB* insertional mutant of high virulent CRAB A0062 (HN510) was constructed as follows: 500 bp of *pmrB* gene was amplified and inserted into psuiAb plasmid that has been digested with HindIII and NcoI (New England Biolabs) using NEB Builder Hifi Assembly (New England Biolabs). The construct was confirmed by PCR and sequencing prior to transformation into A0062. Successful transformants were selected on LB agar plates with apramycin.

### Construction of chromosomal mutants in Salmonella enterica serovar Typhimurium

Chromosomal mutants were constructed with the one-step disruption method with minor modifications (Datsenko & Wanner, 2000). To construct *pmrA* (HN53), *pmrB* (HN49), *pmrApmrB* (HN57), or *oxyR* (BR11) mutant strain, a chloramphenicol cassette was amplified from plasmid pKD3, then introduced into wild-type *Salmonella* 14028s harboring plasmid pKD46. The resulting strains were kept at 30°C and transformed with pCP20 to remove the cat cassette. All mutant strains were confirmed by PCR and sequencing.

### Construction of substitution for amino acid residues via site-directed mutagenesis

The substitution of amino acids was carried out via the site-directed mutagenesis for large plasmids (SMLP) (Zhang *et al*, 2021). SMLP relies on a specialized DNA polymerase to generate two large DNA fragments with overlapping ends by two independent PCRs, using two pairs of partially complementary primers (Appendix Table S2) that contain the substitution mutation. The first DNA fragment (∼500 bp) was amplified via PCR using Q5 High Fidelity 2X Master Mix (New England Biolabs), pWH1266-*pmrB* as template and primers MAFP/MRP. The second DNA fragment (∼ 10,000 bp) was amplified using primers MFP/MARP. Two DNA fragments were joined using NEB Builder Hifi Assembly at 50°C for 15 min. The resulting assembly was transformed into chemically compenent DH5α by heat shock, and successful transformants were selected on LB agar plates with ampicillin following overnight growth at 37°C. The constructs were confirmed by PCR and sequencing prior to introduction to *pmrB* mutant.

### Bacterial growth assays

Freshly streaked bacterial strains were cultured overnight in LB broth at 37°C under shaking. Overnight cultures were sub-cultured at an OD_600_ of 0.005 in LB both without or with drugs at the concentrations indicated in figures and incubated at 37°C with shaking for 20 h. The drugs include H_2_O_2_ (Sigma-Aldrich)_;_ 2,2’- bipyridyl (Sigma-Aldrich); deferoxamine mesylate salt (Sigma-Aldrich), and thiourea (Sigma-Aldrich). Growth was measured by OD_600_ after 20 h of incubation. Growth rate was calculated by dividing the value of OD_600_ of drug-treated bacteria by the value of OD_600_ of non-treated bacteria

### Bacterial survival assays

Freshly streaked bacterial strains were grown overnight in LB broth at 37°C under shaking. Overnight cultures were sub-cultured at an OD_600_ of 0.005 in LB broth and grown to early-stationary phase (6 h). Bacteria were diluted to an OD_600_ of 0.5, washed twice with phosphate buffered saline (PBS, pH 7.4) (137 mM NaCl, 2.7 mM KCl, 10 mM Na_2_HPO_4_, and 1.8 mM KH_2_PO_4_). Then bacteria were re-suspended in PBS without or with drugs and incubated at 37°C with shaking for 2 h. The drugs include H_2_O_2_ , colistin (Sigma-Aldrich), LL-37 (Sigma), deferoxamine mesylate salt, streptonigrin from *Streptomyces flocculus* (Sigma-Aldrich), and thiourea. Name of the drug and its concentration are indicated in figures. The live bacterial numbers or colony-forming units (CFUs) were determined by serial dilution and plating. Bacterial survival was calculated by dividing the number of CFU of drug-treated bacteria by the number of CFU of non-treated bacteria.

For experiments to investigate priming effect on bacterial survival, a similar protocol was used. However, bacteria were primed with 5 µM H_2_O_2_ in PBS for 1 h before being treated with high concentrations of the drugs.

### Reporter assays

To construct reporter plasmids (pLPV1Z-P*_hscB-hscA-fdx_*, pLPV1Z-P*_ahpF1_*, pLPV1Z-P*_pmrC_*, pLPV1Z-P*_naxD_*, pLPV1Z-P*_ftnA_*, pLPV1Z-P*_katE_*, and pLPV1Z-P*_katG_*), promoter regions of the *hscB-hscA-fdx* operon, *ahpF1*, and the *pmrC-pmrA-pmrB* operon, *naxD*, *ftnA*, *katE*, and *katG,* respectively, were amplified using the primers listed in Appendix Table S3 (Lucidi *et al*, 2019). pLPV1Z was digested with PstI and XbaI (New England Biolabs). The purified PCR product and cut plasmid were joined using NEB Builder Hifi Assembly (New England Biolabs) at 50°C for 15 min. The resulting assembly was transformed into chemically compenent DH5α by heat shock, and successful transformants were selected on LB agar plates with gentamicin following overnight growth at 37°C. The constructs were confirmed by PCR and sequencing prior to introduction of this vector to wild-type and mutants. Successful transformants were selected on LB agar plates with gentamicin. For experiments to check gene expression (Fig. 2B and E; Appendix Fig. S4B, E, and G; and Appendix Fig. S12A-B), bacterial strains harboring pLPV1Z-P*_hscB-hscA-fdx_*, pLPV1Z-P*_ahpF1_*, pLPV1Z-P*_pmrC_* or pLPV1Z-P*_naxD_*were grown overnight in LB broth at 37°C under shaking. Overnight cultures were sub-cultured at an OD_600_ of 0.005 in LB broth and grown to early-stationary phase (6 h). 100 µL of bacterial culture was added to 100 µL of LB broth without and with drugs in 96-well white plate. Name of the drug and its concentration are indicated in figures. The luminescence and OD_600_ value were recorded using the BioTek Synergy H1 plate reader. For experiments with metal ions (Fig. 4A-B; and Appendix Fig. S9D-F), the procedure is similar. However, M9 medium was used instead of LB broth.

For priming experiments (Fig. 6B-D; Appendix Fig. S12A and B; and Appendix Fig. S13A-C), bacterial strains with the reporter plasmids were grown overnight in LB broth at 37°C under shaking, subcultured at an OD_600_ of 0.005 in LB broth for 5 h and treated with different concentrations of H_2_O_2_ (1 µM, 5 µM, and 10 µM), as indicated in figures, for 1 h. Then 100 µL of primed bacterial culture was added to 100 µL of LB broth without and with high concentration of drugs in 96-well white plate. Name of the drugs and its concentration are indicated in figures. The luminescence and OD_600_ value were recorded using the BioTek Synergy H1 plate reader.

For experiments to examine how long the priming effect remains effective (Fig. 7B-D), the procedure is similar to the priming experiments. However, primed bacteria were washed twice to remove the prime signal (H_2_O_2_), re-suspended in warm LB broth, and rest for different times (0, 30, 60, 90, and 120 min) before being treated with high concentration of drugs. The luminescence and OD_600_ value were recorded using the BioTek Synergy H1 plate reader.

### PmrA protein purification

Full-length PmrA of *A. baumannii* was obtained though PCR and inserted into the pET28a (+) vector with 6x His tag at the N-terminus. pET28a-PmrA was transformed into BL21 (DE3) *E. coli* competent cells. The cells were grown overnight in LB broth at 37°C with shaking. Overnight culture was sub-cultured 1:100 in LB broth and grown at 37°C with shaking to an OD_600_ of 0.4. Culture was then induced with 1 mM of isopropyl β–D-1-thiogalactopyranoside (IPTG, Sigma-Aldrich) for 3 h at 37°C with shaking. Cells were harvested and pellet was resuspended in binding buffer (pH 7.4) containing tris buffered saline (TBS, Biosesang) and 20 mM of Imidazole (Sigma-Aldrich). Cells were lysed by sonication, and the resulting clarified supernatant was loaded onto a Econo-Pac Chromatography Column (Bio-Rad) which has been pre-loaded with Nickel Sepharose High Performance (Cytiva) and equilibrated with binding buffer. After overnight incubation at 4°C with rotating, the column was washed three times with wash buffer (pH 7.4) containing TBS and 80 mM Imidazole. PmrA protein was eluted using elution buffer (pH 7.4) containing TBS and 250 mM Imidazole. Fractions containing pure protein were then collected after SDS-PAGE, combined and concentrated using Amicon Ultra centrifugal filters (Merck Millipore). Protein concentration was determined using BCA Protein Assay kit (Pierce) according to the manufacturer’s protocol.

### Electrophoretic mobility shift assay (EMSA)

EMSA assays were performed using the LightShift Chemiluminescent EMSA kit (Thermo Scientific). First, 16 µg of PmrA protein was phosphorylated by addition of 7 mM MgCl_2_ (Sigma-Aldrich) and 35 mM BeF_3_^−^ (Sigma-Aldrich) and incubated in room temperature for 1 h. Biotinylated DNA fragments from the upstream regions of *hscB-hscA-fdx* and *ahpF1* were obtained by PCR using 5’-end biotin-labeled primers listed in Appendix Table S3. Binding reactions were incubated in 20 µL volume containing 16 fmol of DNA and various concentrations of phosphorylated PmrA protein, in 1x Gel Shift Binding Buffer (Molecular Depot) for 30 min. Binding reactions were loaded with the provided loading dye onto a 6% Novex TBE Gel (Invitrogen) and separated in 0.5x Novex TBE Running Buffer (Thermo Fisher) at 80 V for 90 min. A Trans-blot Turbo Transfer System (Bio-Rad) was used to transfer proteins and DNA to Biodyne B positively charged nylon membranes (Thermo Fisher) for 30 min at 25 V. Cross-liking was performed by exposing the membranes to UV light for 15 min in a Dual LED Blue/White Light Transilluminator (Invitrogen). Blocking, incubation with streptavidin-horseradish peroxidase conjugate, washing and revealing were performed according to the protocol. Membranes were imaged using Amersham ImageQuant 900 (Cytiva).

### Subcellular Localization of the PmrB Proteins

Isolating bacterial inner membrane (IM) from outer membrane (OM) and cytosol (Cy) of *A. baumannii* was performed using a protocol described previously (Cian *et al*., 2020). Briefly, *pmrB* mutant harboring pWH1266-*pmrB*-3xFLAG or pWH1266-*pmrB1*-3xFLAG was grown overnight in LB medium at 37°C under shaking. Overnight cultures were sub-cultured at an OD_600_ of 0.005 in 1 liter of LB medium and grown to early-stationary phase (6 h). Cells were harvested by centrifugation at 7,000 x g for 10 min at 4°C. Cells were re-suspended in 12.5 mL of buffer A (0.5 M sucrose, 10 mM Tris pH 7.5, 144 µg/mL lysozyme) and 12.5 mL of 1.5 mM EDTA. Cell pellet was collected by centrifugation at 7,000 x g 10 min at 4°C and re-suspended in 25 mL buffer B (0.2 M sucrose, 10 mM Tris pH 7.5, 1 M MgCl_2_, 5 U/mL Dnase I, one tablet of protease inhibitor cocktail in 25 mL of solution). Cells were then disrupted by sonication and cell debris was removed by centrifugation at 7,000 x g for 1 h at 4°C. Total membrane fraction was collected by centrifugation of the the whole-cell lysate at 184,500 x g for 1 h at 4°C. The total membrane fraction was re-suspended in 1 mL of low-density isopycnic sucrose gradient solution (20% w/v sucrose, 1 mM EDTA, 1 mM Tris Ph 7.5), loaded on top of a sucrose gradient made with 4 mL of medium-density isopycnic sucrose gradient solution (53% w/v sucrose, 1 mM EDTA, 1 mM Tris Ph 7.5) and 2 mL of high-density isopycnic sucrose gradient solution (73% w/v sucrose, 1 mM EDTA, 1 mM Tris Ph 7.5) in a Beckman Ultra-Clear centrifuge tube followed by centrifugation in an SW41 rotor at 288,000 x g at 4°C for 20 h. Bands between 20% and 45% (upper, reddish band) and between 45% and 73% (lower, white band), corresponding to the inner and outer membranes, respectively, were collected. Isolated membranes were washed and stored with membrane-storage buffer (10 mM Tris Buffer pH 7.5).

### Western blot analysis

Protein concentrations were determined using BCA Protein Assay Kit (Pierce), with bovine serum albumin (BSA) used as a standard protein. Protein samples (100 µg of protein each) were run in a Bolt 4-12% Bis-Tris Plus gel (Invitrogen) and transferred onto a nitrocellulose membrane using iBlot 2 Gel Transfer Device (Invitrogen). Membranes were blocked with 5 % skim milk solution at room temperature for 1 h. Then, samples were analyzed using anti-FLAG antibody (Sigma-Aldrich) (1:2,000 dilution) and secondary horseradish peroxidase-linked antibody (Cytiva) (1:5,000). The blots were developed with ECL detection system (Thermo Scientific). Membranes were imaged using Amersham ImageQuant 900 (Cytiva).

### Β-Nicotinamide adenine dinucleotide (NADH) oxidation assays

To confirm inner membrane (IM) fractions, enzymatic activity of NADH dehydrogenase, which exists exclusively in the IM, was measured. 50 µg of fractioned protein was added to 10 mM Tris-buffer (pH 8.0) to make up a final volume of 990 µL. Then 10 µL of a 10 mgmL solution of NADH (Roche) was added to the sample and absorbance at 340 nM was measure every 5 min for 10 h using using the BioTek Synergy H1 plate reader.

### Quantitative RT-PCR

To measure the relative mRNA levels, bacteria were grown in LB broth at 37°C under shaking. Overnight cultures were sub-cultured at an OD_600_ of 0.005 in LB broth and grown to early-stationary phase (6 h). 750 µL of bacterial culture was added to 750 µL pf RNAprotect Bacteria Reagent (QIAGEN). Bacterial pellets were collected by centrifugation at 13,000 rpm for 10 min at 4°C. Total RNA was purified by using Rneasy Mini Kit (QIAGEN) with on-column DNAse treatment, and cDNA was synthesized by using iScript Reverse Transcription Supermix (Bio-rad). Quantification of transcripts was carried out by qRT-PCR using Fast SYBR Green Master Mix (Thermo Fisher) in an CFX384 Real-Time PCR Detection System (Bio-rad). The relative amount of mRNA was determined by using a standard curve obtained from PCR with serial diluted genomic DNA, and results were normalized to the levels of 16S rRNA. The mRNA levels of genes were measured by sing the indicated primer sets (Appendix Table S3). Data shown are an average from three independent experiments.

### PmrB protein purification

All PmrB proteins used in this study were expressed and purified as previously reported (Bhogaraju *et al*, 2016). *pmrB^30-140^*, *pmrB^30-140(His86,89,90,92Ala)^*were cloned into pParallel GST2 vector. BL21(DE3) *Escherichia coli* competent cells (NEB) were transformed with *pmrB* plasmids and grown in LB medium to an OD_600_ of 0.6 at 37°C. Protein expression was then induced by the addition of 0.5 mM IPTG for 16 h at 17°C, after which cells were harvested. The cell pellet was resuspended in lysis buffer (50 mM Tris-HCl [pH 8.0], 150 mM NaCl, 1 mM PMSF), lysed by sonication, and centrifuged at 13,000 rpm for 1 h to collect the supernatant. The supernatant of GST-tagged proteins was incubated for 2 h with glutathione Sepharose 4B (Cytiva), pre-equilibrated with wash buffer (50 mM Tris-HCl [pH 8.0], 500 mM NaCl), for cleared non-specifically bound proteins. Glutathione beads were incubated with sfGFP-TEV protease for 2 h at 25°C to cleave the GST-tag. The cleaved proteins were exchanged into IEX buffer A (20 mM Tris-HCl [pH 8.0], 20 mM NaCl) and purified by anion-exchange chromatography using HitrapQ column (Cytiva), with gradient elution using IEX buffer B (20 mM Tris-HCl [pH 8.0], 1 M NaCl). Purified proteins were loaded onto size-exclusion column (Superdex 75 16/60, Cytiva), and pre-equilibrated with 20 mM Tris-HCl [pH 8.0], 20 mM NaCl.

### Inductively Coupled Plasma-Mass Spectrometry (ICP-MS) analysis

The metal element and quantity of PmrB proteins were determined by ICP-MS (NexION 350D, Perkin-Elmer SCIEX) using argon plasma (6,000K) and high mass resolution (0.3-3.0amu). For sample preparation, the protein was extracted from whole cells and purified using size exclusion chromatography (Superdex 75 16/60, Cytiva) using 20 mM Tris-HCl [pH 8.0], 20 mM NaCl buffer. And then, samples are injected by 1.00 mL/min flow rate and measure the concentration of metals (Mn^2+^, Co^2+^, Ni^2+^ and total Fe).

### Protein cofactor analysis by fluorescence dye

Protein samples were denatured using 6M guanidine hydrochloride (pH 8.5) (Sigma-Aldrich) and then incubated with 10 µM Newport Green (Invitrogen) for 60 min at room temperature prior to analysis. For control, BSA (Sigma-Aldrich) was used. Fluorescence was recorded using the BioTek Synergy H1 plate reader set at an excitation 485 nM and an emission of 538 nM.

### Intramacrophage survival Assay

For the intracellular survival assays, J774A.1 cells were seeded in 24-well plates and incubated at 37°C and 5% CO_2_ for 36 h before the experiment (2×10^4^ cells/well). *A. baumannii* strains were grown overnight in LB at 37°C under shaking condition. Overnight cultures were sub-cultured at an OD_600_ of 0.005 in LB and grown to early-stationary phase (6 h). An appropriate volume was added to each well to reach an MOI of 100. Plates were centrifuged 10 min at 700 x g to enhance bacterial contact with host cells and incubated for 1 h at 37°C and 5% CO2. Cells were washed twice with PBS and subsequently incubated with medium supplemented with 100 µg/mL gentamicin to kill extracellular bacteria. At 2 h post infection (p.i.), the concentration of gentamicin was decreased to 10 µg/mL. At 2 h and 6 h p.i., cells were washed twice with PBS and lysed with 500 µL PBS containing 0.2% Triton X-100. Lysates were serially diluted and plated on LB agar plates to determine CFU. Percentage of bacterial survival was calculated by dividing the number of CFU at 6 h p.i. by the number of CFU at 2 h p.i.

### Lung infection of A. baumannii strains in mice

Acute lung infection was induced by intratracheal instillation of *A. baumannii* in wild-type (WT) mice (C57B6/J, Jax:00664) and R26LSL-Tomato mice (Jax: 007909) while mice were lightly sedated with isoflurane. Mice received treatment with either 20 µL of sterile saline or *A. baumannii* (10^6^ CFU).

### Tissue preparation and single-cell suspension

To analyze blood leukocytes, blood was obtained by needle puncture from submandibular vessels. Red blood cells (RBC) were removed by blood lysis buffer (BD). The entire lung tissue was minced and enzymatically digested in PBS containing collagenase type IV (Sigma-Aldrich, 1 mg/mL), hyaluronidase (Sigma-Aldrich, 0.15 mg/mL), and Dnase I (Roche. 0.05 mg/mL) at 37 °C with agitation (250 rpm) for 30 minutes. After tissue digestion, cell suspensions were filtered through a 70 µm cell strainer (Corning), then total cells were counted by an automated cell counter (Nexcelom). To quantify neutrophils in the lungs, the total cell count was multiplied by the percentages of CD45+ CD11b+Ly6G+ neutrophils.

### Flow cytometric analysi

Single-cell suspensions collected from the lung tissue were maintained on ice for flow cytometry antibody staining. Antibodies targeting CD45 (30-F11), Ly6C (HK1.4), CD115 (AFS98), Ly6G (1A8), and CD11b (M1/70) were used to label leukocytes in blood and lungs for flow cytometric analysis. The labeled cell suspension was analyzed by LSR Fortessa instrument (BD Biosciences). The acquired data were further analyzed the FlowJo software v10.9 (Tree Star).

### H&E Staining

Histological staining was performed on paraffin sections using standard methods. Tissues were fixed overnight in 10% formalin at 4°C, followed by dehydration, embedded in paraffin, and slicing into 10 μm-thick sections using a microtome. The tissues were deparaffinized, and endogenous peroxides were blocked with a 3% H_2_O_2_ solution. The sections were stained with hematoxylin and eosin (H&E). Sample images were captured using a Leica Dmi8 microscope (Leica). Quantitative analysis of H&E staining was performed using ImageJ. Immune cell–dense regions were identified by gray-scale thresholding of hematoxylin/eosin-positive tissue. Tissue-positive area and total image area were measured from the thresholded masks, and immune cell area (%) was calculated as tissue-positive area divided by total area.

### Immunohistochemistry (IHC) analysis

The lungs obtained from Ly6gCre:R26LSL-Tomato mice were initially fixed overnight in 4% paraformaldehyde (Thermo Fisher Scientific) at 4°C, then lung tissues were cryoprotected in 30% sucrose at 4°C for 2-3 days. Subsequently, lung tissues were embedded in optimal cutting temperature compound (OCT, Sakura Finetek, USA). The tissues were snap-frozen in liquid nitrogen, followed by storage at -80°C. The frozen tissues were sectioned at a thickness of 10 µm with a cryostat (Leica, CM1800). Nuclei were stained with bis Benzimide H 3342 (Sigma-Aldrich). The lung sections were imaged by Zeiss 880 confocal microscope. The acquired images were processed with Imaris software (Bitplane).

### Orthologous gene identification and acquisition for PmrB

We utilized the OrthoDB database to investigate the orthologous genes of *A. baumannii* PmrB (444 amino acids) (Kuznetsov *et al*, 2023). The input ID “470_0:000b47” provided us with the group 9809766at2 at the bacteria level, yielding a FASTA file with 8,597 genes across 3,981 species.

### Data preparation for PmrB comparison analysis

Using an in-house Python script, the dataset was renamed to facilitate compatibility with high-throughput analysis tools. The sequences were then aligned using MAFFT version 7 to generate a Newick tree file (Katoh *et al*, 2019). We selected the 400 genes most closely related to our target gene for further analysis. In instances where multiple strains of the same species were present, only the closest relative to the target gene was retained. Additionally, *Salmonella enterica* was included as a control.

### Multiple sequence alignment and visualization for Pmr

For the multiple sequence alignment, we adopted a two-tiered approach. Representative species from each genus were randomly selected for an initial alignment to encapsulate broad genetic diversity. In contrast, within the genus *Acinetobacter*, which includes our target organism, all species were aligned separately to provide a detailed view of the genetic variation within this specific group. We utilized Clustal Omega for the alignment processes, ensuring accurate representation of the evolutionary relationships. The visualization of the aligned sequences was performed using BioEdit version 7.7.1 which allowed for the generation of illustrative graphics depicting the comparative sequence homology (Hall, 1999).

### Phylogenetic tree construction and representation for PmrB

The phylogenetic tree was constructed using Clustal Omega, based on the curated FASTA file. We visualized the tree using iTOL version 6.8.1, color-coding the terminal branches to represent different genera and creating an outer donut-shaped chart that represents familial and class levels of taxonomy for additional context (Ciccarelli *et al*, 2006).

### Gene presence estimation strategy for rpoS and iron-sulfur assemble systems

In this study, we implemented a strategic approach to estimate the presence of genes across various genera. For genera that were included in the phylogenetic analysis, we randomly selected one species from those used in the analysis. For genera not included in the phylogenetic analysis, we randomly chose one species from the available options. This methodology was focused on accurately assessing the number of registered genes per species while eliminating duplicates due to isoform presence.

Using the orthologous group information from OrthoDB for genes such as *rpoS* (9809557at2), *nifA* (9811901at2), *nifB* (53113at2157), *nifD* (9762718at2), *39nife* (9767044at2), *nifF* (359268at2), *nifH* (9778641at2), *nifJ* (9794954at2), *nifK* (9800746at2), *nifL* (8579121at2), *nifN* (134630at2157), *nifQ* (192277at2), *nifS* (9808002at2), *nifU* (319865at215), *nifV* (503431at2), *nifW* (9811868at2), *nifX* (9797941at2), *sufA* (52656at2157), *sufB* (9803529at2), *sufC* (9806149at2), *sufD* (9768262at2), *sufE* (9799320at2), *sufS* (9808002at2), and *sufU* (9804157at2), we calculated the gene counts for each species. This allowed us to estimate the presence of the *rpoS*, *nif*, and *suf* systems within the genomes under investigation. In particular, considering the complexity of the *rpoS* (9809557at2) orthologous group, which includes a significant number of paralogs, our analysis was refined to utilize only data accurately registered as *rpoS*.

### Quantification and statistical analysis

Data and statistical analysis were performed using GraphPad Prism 10.2.2. Replicates and statistical details can also be found in the methods and figure legends.

### Protein design and structure prediction

A PmrB periplasmic domain (amino acid residues 29-136) was designed as a Ni^2+^-responsive reactive oxygen species (ROS) sensor, incorporating a histidine-rich metal-binding motif (HLSHHLH) centered at residues 86–92. The binding site comprises four histidine residues (His86, His89, His90, and His92) arranged to coordinate Ni^2+^ ions in a square-planar-like geometry amenable to redox-dependent conformational changes. The three-dimensional structure was predicted using ESMFold (Lin *et al*, 2023), a protein language model-based structure prediction method. The predicted structure served as the initial conformation for all subsequent molecular dynamics simulations.

### System Preparation and Bonded Model

Three simulation systems were prepared: (i) Apo (metal-free), (ii) Ni^2+^-bound (reduced), and (iii) Ni^3+^-bound (oxidized). The protein was parameterized using the AMBER ff14SB force field (PMID: 26574453) with TIP3P-FB water (PMID: 28306259). Each system was solvated in a periodic box with a minimum padding of 1.0 nm from the protein to the box edge, and Na^+^/Cl^−^ counterions were added to achieve a physiological ionic strength of 0.15 M.

For the Ni^2+^- and Ni^3+^-bound states, bonded model parameters for the metal–ligand interactions were derived using the Seminario method (Seminario, 1996) applied to Hessian matrices computed at the GFN2-xTB level of theory (Bannwarth *et al*., 2019). A Ni–His_4_ cluster comprising the four coordinating histidine sidechains and the nickel ion was extracted from the predicted structure. Geometry optimization and frequency calculations were performed using xtb (version 6.6) for both the Ni^2+^ (charge = +2) and Ni^3+^ (charge = +3) oxidation states. Bond stretching (Ni–Nε) and angle bending (N–Ni–N) force constants were extracted from the projected Hessian matrices following the Seminario approach, yielding equilibrium bond lengths and force constants specific to each oxidation state. The resulting parameters were formatted as AMBER frcmod files and incorporated into the system topology via tleap (AmberTools 23) (Case *et al*, 2023). Explicit Ni–Nε bonds were defined in tleap to enforce the coordination geometry throughout the simulations.

### Molecular Dynamics Simulations

All MD simulations were performed using OpenMM 8.4 (Eastman *et al*., 2024) on an NVIDIA A100 GPU with CUDA mixed-precision arithmetic. Each system was subjected to energy minimization followed by sequential equilibration: 100 ps under the NVT ensemble and 100 ps under the NPT ensemble with harmonic restraints on heavy atoms that were gradually released. Production simulations were carried out for 100 ns under the NPT ensemble at 300 K using the Langevin Middle integrator (γ = 1.0 ps^−1^) and 1 atm using the Monte Carlo barostat. A 2 fs integration time step was used with the HBonds constraint on all bonds involving hydrogen atoms. Hydrogen mass repartitioning (HMR, 1.5 amu) was applied (Hopkins *et al*, 2015). Non-bonded interactions were treated with a 1.0 nm cutoff, and long-range electrostatics were computed using the particle mesh Ewald (PME) method (Hess *et al*, 2008). Trajectory frames were saved every 10 ps, yielding 10,000 frames per trajectory. Three independent replicates (n = 3) with different random velocity seeds were performed for each system, yielding a total of 900 ns of aggregate simulation time.

### Structural and Dynamic Analysis

All trajectory analyses were performed using MDTraj 1.9 (McGibbon *et al*., 2015) and MDAnalysis 2.7 (Michaud-Agrawal *et al*., 2011). Root-mean-square deviation (RMSD) was calculated for all Cα atoms and for the binding site residues (His86, His89, His90, and His92) relative to the initial minimized structure. Per-residue root-mean-square fluctuation (RMSF) was computed after least-squares superposition. Differential RMSF (ΔRMSF = Ni^3+^ − Ni^2+^) was calculated with propagated uncertainties (σ_Δ_ = √(σ_Ni3+_^2^ + σ_Ni2+_^2^)).

Principal component analysis (PCA) was performed on Cα coordinates using scikit-learn (Pedregosa *et al*, 2011). Each trajectory was superposed onto the first frame, and projections onto PC1 and PC2 were visualized with time-dependent coloring (viridis colormap). Solvent-accessible surface area (SASA) was computed using the Shrake–Rupley algorithm (Shrake & Rupley, 1973) as implemented in MDTraj. Radius of gyration (R_g_) was calculated to evaluate protein compactness. Differential contact maps (ΔContact) were generated from the difference in average inter-residue Cα–Cα distance matrices between Ni^3+^ and Ni^2+^ states using the last 50 ns, averaged across three replicates. Ni–His Nε coordination distances were monitored using MDAnalysis. All results are presented as mean ± SD (n = 3). Figures were generated using Matplotlib 3.8 (Hunter, 2007).

## Data availability

All data are available in the main text or the supplementary materials. All simulation and analysis code is deposited at GitHub (https://doi.org/10.5281/zenodo.18780313).

## Author Contributions

H.V.N., S.H.K., and J.Y. conceived and designed the experiment. H.V.N., S.H.K., H.H., S.K., D.S., K.K., and J.Y. conducted experiments. M.G. provided materials. H.V.N., S.H.K., K.K., and J.Y. wrote the manuscript. All authors reviewed and approved the manuscript.

## Acknowledgments

We thank Prof. Eduardo Groisman at the Yale University for comments on the early draft of manuscript. We are also grateful to Prof. Bryan Davies at the University of Texas at Austin for providing pAT02 and pAT03 plasmids, and Prof. Paolo Visca at the Roma Tre University for providing pLPV1Z plasmid. This work was supported by the New Faculty Startup Fund from Seoul National University to J.Y., the National Research Foundation Korea Basic Science Research Programs (2021R1C1C1005184, RS-2026-25489275 and 2020R1A5A1019023 to J.Y.), the Global-LAMP Program of the National Research Foundation of Korea (NRF) grant funded by the Ministry of Education (RS-2023-00301976), the Korea Health Technology R&D Project through the Korea Health Industry Development Institute (KHIDI) (HI23C026400 and RS-2023-00304637), and Brain Pool program by the National Research Foundation of Korea (2021H1D3A2A02083176 to H.V.N).

